# Epigenetic regulation of DPP4 receptor expression by NONO enables replication of MERS-CoV

**DOI:** 10.64898/2026.09.21.753343

**Authors:** Adam Hage, Mikhaila Janes, Seth D. Scott, Suhyeon Yoon, Kevin Rose, Tovah E. Markowitz, Lara M. Myers, Aaron B. Carmody, Lenny Triem, Olivia Straw, Kristin L. McNally, Paul A. Beare, Justin B. Lack, Craig Martens, Sonja M. Best

## Abstract

Middle East respiratory syndrome coronavirus (MERS-CoV), first reported in 2012, belongs to the *Betacoronavirus* genus, including SARS-CoV and SARS-CoV-2. Human-to-human transmission of MERS-CoV appears inefficient, but repeated spillover events have been reported from at least 27 countries, raising concern for future emergent events. Coronaviruses (CoVs) rely extensively on host proteins to support their replication. Several studies have implicated the paraspeckle non-POU domain-containing octamer-binding protein (NONO) as an RNA-binding protein (RBP) that binds to CoV genomes, but the role of this protein in regulating replication is unknown. Here, we show that NONO is required for expression of dipeptidyl peptidase 4 (DPP4, also known as CD26), the major cellular receptor required for MERS-CoV attachment and entry. NONO did not impact *DPP4* mRNA processing or stability but instead promoted *DPP4* transcription through control of active H3K4me3 and repressive H3K27me3 histone modifications at the *DPP4* locus. Together, these findings identify NONO as a key proviral host factor for MERS-CoV and reveal an epigenetic mechanism linking a host RBP to viral entry receptor expression that may represent a target for therapeutic strategies.

## Introduction

Within the last 25 years, the world has experienced the emergence of three highly pathogenic coronaviruses (CoVs); severe acute respiratory syndrome coronavirus (SARS-CoV) in 2002, Middle East respiratory syndrome coronavirus (MERS-CoV) in 2012, and the recent pandemic severe acute respiratory syndrome coronavirus 2 (SARS-CoV-2) in 2019. SARS-CoV-2 is responsible for a currently estimated 7.1 million cumulative global deaths as tracked by the World Health Organization (WHO). Human-to-human transmission of MERS-CoV appears inefficient, but repeated spillover events have been reported from at least 27 countries, raising concern for future emergent events. Emerging respiratory viruses pose an important public health threat and an incomplete understanding of the molecular determinants of viral pathogenesis can become an important barrier to the development of treatment options.

Viral infection of a host cell is a multistep procedure involving attachment and entry, initial transcription of the viral genome and translation of viral proteins, and assembly and egress of nascent virions. Throughout this process, viruses must counteract host antiviral responses that have evolved to impede every step of replication. Despite having a roughly 30 kb size genome, CoVs produce only a few dozen of their own proteins, many of which retain multiple host antagonism functions to hinder antiviral innate immune responses. Due to this limited genome size, viruses often do not encode a sufficient proteome to respond to host countermeasures while also completing viral replication. To overcome this, viruses often “hijack” host proteins involved in the cell cycle and repurpose them as their own machinery. Host RNA-binding proteins (RBPs) regulate RNA processing, splicing, and translation making them attractive targets for viruses. For coronaviruses, multiple studies have been published identifying RBPs bound to viral RNA (vRNA) using comprehensive Identification of RNA-binding proteins by Mass Spectrometry (ChIRP-MS) or RNA antisense purification coupled with mass spectrometry (RAP-MS) in three separate cell lines (HEK293T, VeroE6, and Huh7.5) ^1–3^. Comparisons by Labeau and colleagues have generated a consensus list of 58 host RBPs as high-confidence members of the SARS-CoV-2 vRNA interactome ^3^, although the mechanistic consequences of these interactions are not known. We had interest in one of these proteins, the paraspeckle non-POU domain-containing octamer-binding protein (NONO), as a regulator of host innate immunity ^4^, and therefore chose to pursue mechanistic studies on the role of NONO in CoV replication.

NONO belongs to the Drosophila behavior/human splicing (DBHS) family of proteins which includes splicing factor proline/glutamine rich (SFPQ/PSF) and paraspeckle protein component 1 (PSPC1/PSP1). NONO, SFPQ, and PSPC1 are key RBPs for the formation of paraspeckle nuclear bodies, gene regulatory condensates that are functionally important during cellular stress responses by trapping proteins and mRNAs and enhancing pre-mRNA processing ^5^. DBHS proteins are known to participate in nearly every aspect of host RNA metabolism and recent evidence indicates DBHS proteins may have diverse functions during viral infection including both proviral and antiviral activities. In this study, we establish NONO to be an essential host factor for MERS-CoV replication. Transcriptomic and epigenomic analyses combined with genetic deletion and rescue experiments reveal NONO is required for proper transcription of the cell surface receptor dipeptidyl peptidase 4 (DPP4, also known as CD26). DPP4 is the entry receptor for MERS-CoV and cells lacking NONO failed to express sufficient DPP4 due to a reduction in chromatin accessibility at the DPP4 loci. By integrating meta-analyses from RBP-CoV vRNA interactome studies with Next-generation sequencing and functional characterization, we identify NONO as a MERS-CoV host dependency factor.

## Results

### NONO is an essential host factor for MERS-CoV replication

To determine whether NONO has a role in regulation of coronavirus replication, we first utilized HAP1 cells that are permissive to MERS-CoV replication without the need to overexpress cellular receptors. Wildtype (WT) and *NONO* knockout (KO) HAP1 cells were infected with MERS-CoV at a low multiplicity of infection (MOI) (Figures 1A and 1B). Strikingly, loss of NONO resulted in a 100-to 1000-fold decrease in production of infectious titers (Figure 1C). Interestingly, a large defect in virus production was noted as early as 24 hours post infection (hpi). This impairment in virus replication was evident by immunoblot where accumulation of MERS-CoV nucleocapsid (N) protein was reduced in *NONO* KO cells even at 24 hpi (Figure 1D). To determine if NONO also enhanced SARS-CoV-2 replication, A549 cells stably expressing the human ACE2 and TMPRSS2 entry receptors were treated with a pool of small interfering RNAs (siRNA) to deplete *NONO* expression. Here, we observed a 3-to 4-fold reduction in titers of infectious SARS-CoV-2 beginning at 48 hpi (Figure S1A). Therefore, we focused efforts on further examination of MERS-CoV. To determine if the NONO proviral effect on MERS-CoV replication is direct or indirect, we examined interferon (IFN) and IFN-stimulated gene (ISG) responses in WT and *NONO* KO cells. MERS-CoV infection did not induce significant expression of *IFNB1* and *IFIT1* transcripts, and no differences in gene induction was observed between WT and *NONO* KO cells (Figure 1E) except for a modest increase in *IFNL2* mRNA in MERS-CoV-infected *NONO* KO treatment (2.5-fold increase) (Figure 1E). To rule out IFN-signaling as a possible explanation for the observed phenotype, WT and *NONO* KO cells were treated with the inhibitor Ruxolitinib to prevent JAK phosphorylation and subsequent downstream transcription of ISGs. However, Ruxolitinib treatment did not affect virus titers produced from either cell type, consistent with the minimal induction of IFN-associated gene expression (Figure 1F). Collectively, these data demonstrate that NONO has proviral roles in replication of betacoronaviruses but is critical for MERS-CoV replication independent from modulating the antiviral innate immune response.

**Figure 1.**
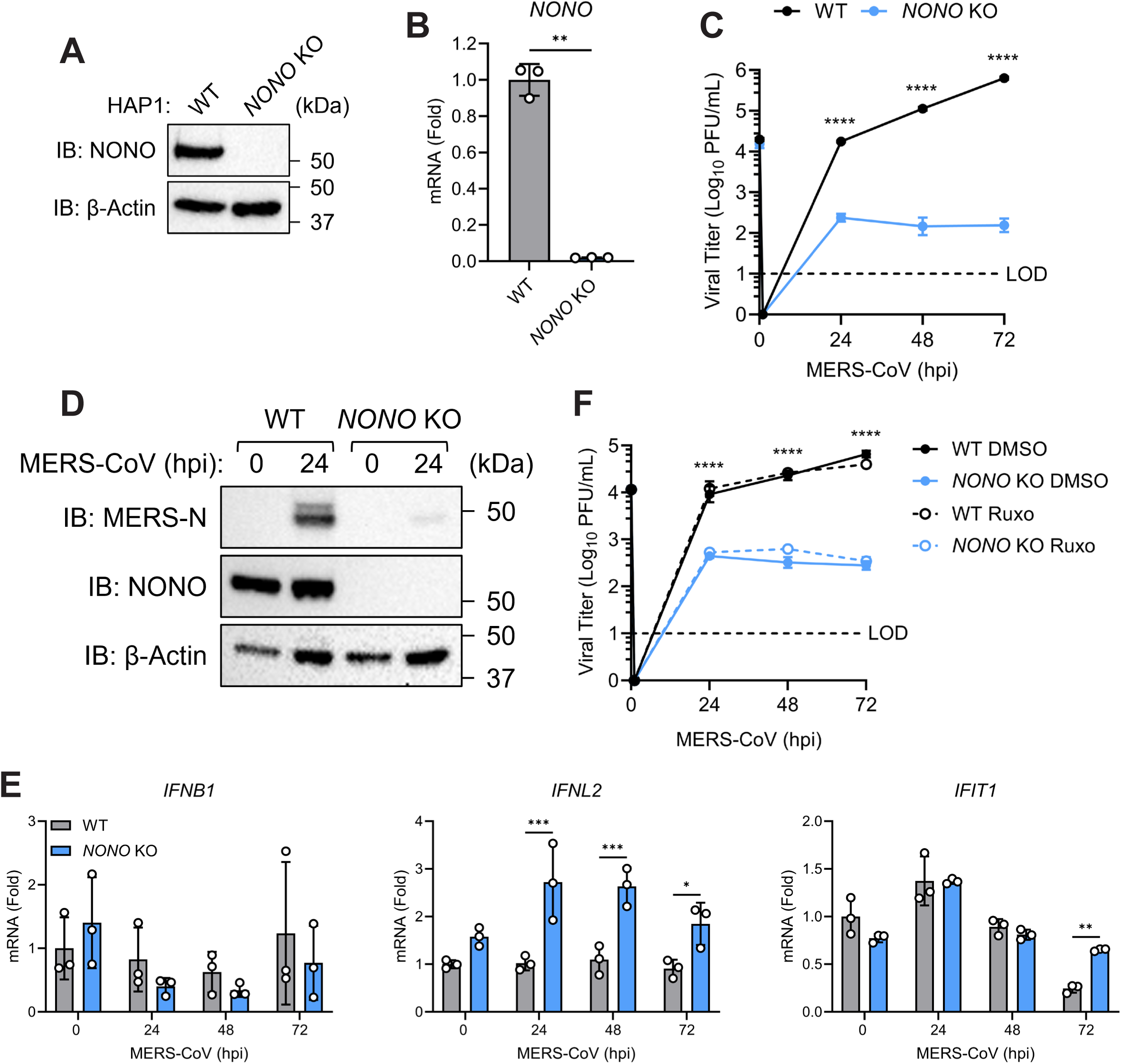
NONO is an essential host factor for MERS-CoV replication: (A-B) HAP1 CRISPR knockout efficiency of (A) NONO protein (immunoblot) and (B) *NONO* mRNA (qRT-PCR). (C-E) WT or *NONO* KO HAP1 cells infected with MERS-CoV (MOI=0.01). (C) Infectious viral titers (plaque assay). (D) Protein levels for MERS-CoV N (immunoblot). (E) *IFNB1*, *IFNL2*, and *IFIT1* expression (qRT-PCR). (F) Infectious viral titers from WT or *NONO* KO HAP1 cells pretreated with Ruxolitinib (10 μg/mL) or DMSO carrier for 1 hour before infection with MERS-CoV (MOI=0.01) (plaque assay). Data are expressed as means (n=3) ± SD, *p < 0.05, **p < 0.01, ***p < 0.001, ****p < 0.0001 (two-way ANOVA with Sidak’s multiple comparisons or Welch’s t-test) and are representative of 2-3 independent experiments.

### NONO supports MERS-CoV vRNA replication and transcription

To determine what stage of the MERS-CoV replication cycle is impacted by NONO, we first examined the consequence of NONO deletion on replication and transcription of the vRNA. Strand-specific qRT-PCR was performed to measure the abundance of MERS-CoV genomic vRNA (*ORF1b* and *upE*) or subgenomic mRNA (membrane (*M*)) transcript species between WT and *NONO* KO cells infected with a low multiplicity of infection (MOI). WT cells supported an increase in abundance of all three MERS-CoV RNAs tested while *NONO* KO cells failed to produce a significant increase over 72 hours of infection (Figure 2A). To understand how early the MERS-CoV replication defect could be observed, we infected cells with MERS-CoV at 4°C for 1 hour before shifting to 37°C. Under these more highly synchronized infection conditions, MERS-CoV failed to replicate genomic RNA or produce *M* mRNA in *NONO* KO cells, which was evident in WT cells as early as 4 hpi (Figure 2B). Thus, NONO has a vital function at an early rate-limiting step in the MERS-CoV replication cycle.

**Figure 2.**
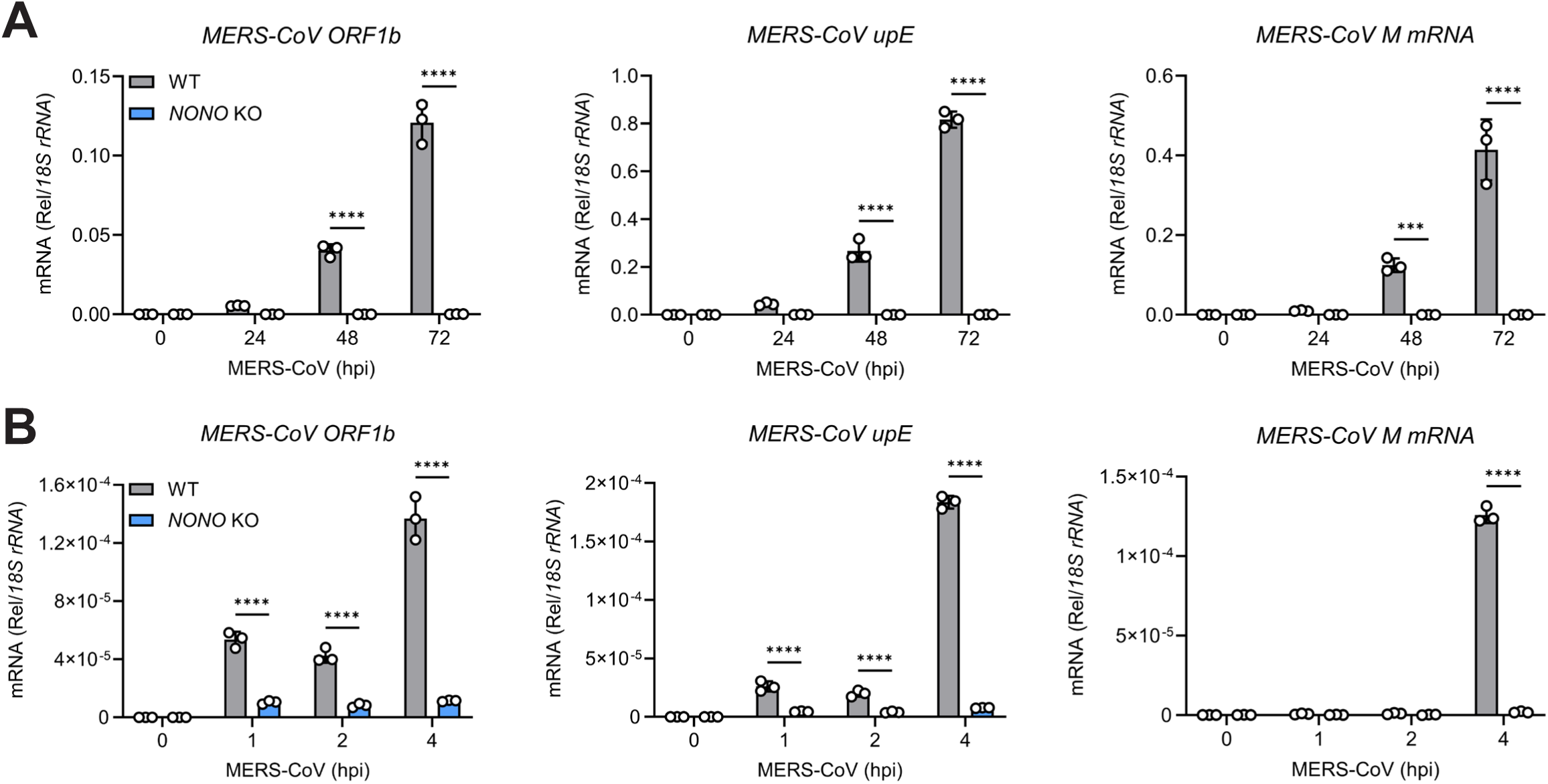
NONO supports MERS-CoV vRNA replication and transcription: (A-B) WT or *NONO* KO HAP1 cells infected with MERS-CoV (MOI=0.01). (A) MERS-CoV *ORF1b*, *upE*, and *M* expression (0-72 hpi) (qRT-PCR). (B) MERS-CoV *ORF1b*, *upE*, and *M* expression (0-4 hpi) from synchronized infection (qRT-PCR). Data are expressed as means (n=3) ± SD, ****p < 0.0001 (two-way ANOVA with Sidak’s multiple comparisons) and are representative of 2-3 independent experiments.

### NONO is required for DPP4 transcription and MERS-CoV entry

The requirement of NONO for MERS-CoV to initiate replication of its vRNA implies that NONO may be needed at the level of viral attachment and entry. To test this, we used a virus-like particle (VLP) system containing all four MERS-CoV structural proteins (S, M, N, and E) that encapsulates an alphavirus-derived defective RNA genome expressing firefly luciferase as a readout for viral entry into target cells. Entry of MERS-CoV VLPs was highly impaired in *NONO* KO cells compared to WT as measured by luciferase levels at 6 hours post incubation (Figure 3A). Therefore, we investigated whether loss of NONO led to altered levels of the primary entry receptor for MERS-CoV, DPP4. *DPP4* mRNA expression was significantly reduced in *NONO* KO cells as observed by qRT-PCR and RNA-seq (Figures 3B and 3C). Gene Set Enrichment Analysis (GSEA) revealed downregulation of mRNA splicing pathways and mRNA 3’-end processing consistent with the role of NONO as a splicing factor (Figure S1B). Upregulated genes in *NONO* KO cells were enriched in pathways identified in our previous NONO investigation in A549 cells (Cite) including eukaryotic and cap-dependent translation initiation among others (Figure S1C). The observed loss of *DPP4* mRNA transcription was consistent with a 2-fold reduction in total protein expression and cell-surface localization (Figures 3D, 3E, S1D, and 3F). We further confirmed that DPP4 is the primary cellular receptor for MERS-CoV entry in HAP1 cells, since knockout of DPP4 expression by CRISPR/Cas9 gene editing completely eliminated MERS-CoV infection without affecting NONO expression (Figures S2A and S2B). These results suggest that NONO is required for optimal expression of DPP4.

**Figure 3.**
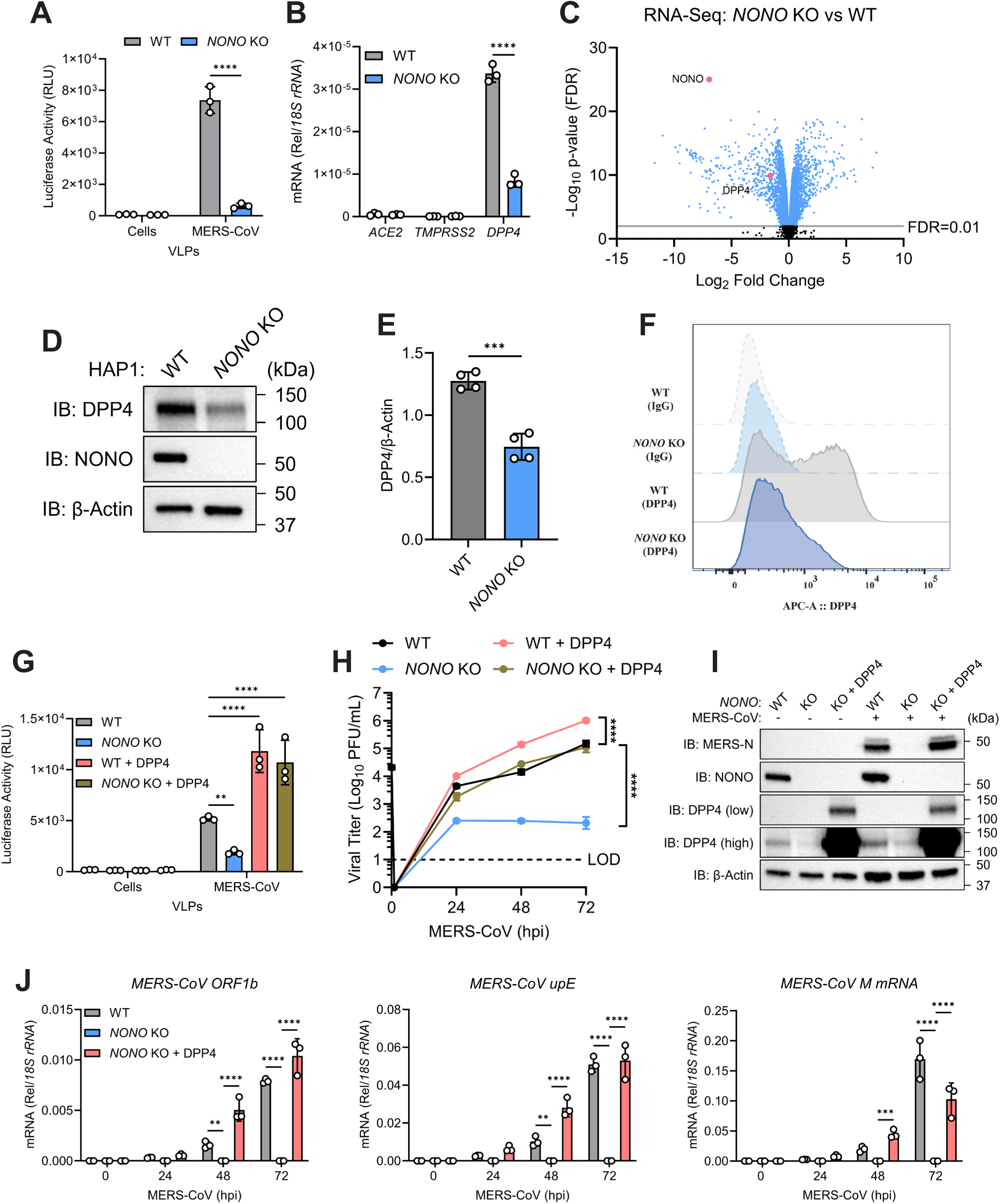
NONO deletion disrupts DPP4 transcription and MERS-CoV entry: (A) Luciferase signal from MERS-CoV VLP entry of WT or *NONO* KO HAP1 cells (VLP luciferase). (B) *ACE2*, *TMPRSS2*, and *DPP4* expression from WT or *NONO* KO HAP1 cells (qRT-PCR). (C) RNA-seq volcano plot showing significant downregulation of DPP4 in *NONO* KO compared to WT samples. (D) Protein levels for DPP4 from WT or *NONO* KO HAP1 cells (immunoblot). (E) DPP4 densitometry analysis from (D). (F) DPP4 surface expression from WT or *NONO* KO HAP1 cells (flow cytometry). (G) Luciferase signal from MERS-CoV VLP entry of WT or *NONO* KO HAP1 cells with or without DPP4 transduction (VLP luciferase). (H-J) WT or *NONO* KO HAP1 cells with or without DPP4 transduction infected with MERS-CoV (MOI=0.01). (H) Infectious viral titers (plaque assay). (I) Protein levels for MERS-CoV N and DPP4 (immunoblot). (J) MERS-CoV *ORF1b*, *upE*, and *M* expression (qRT-PCR). Data are expressed as means (n=3) ± SD, **p < 0.01, ***p < 0.001, ****p < 0.0001 (two-way ANOVA with either Sidak’s, Tukey’s, or Dunnett’s multiple comparisons or Welch’s t-test) and are representative of 2-3 independent experiments.

To confirm that loss of DPP4 is the primary reason for the MERS-CoV replication disadvantage in *NONO* KO cells, we transduced DPP4 into WT and *NONO* KO cells using lentiviruses to generate stable cell lines with restored surface level expression of DPP4 (Figure S2C). Entry of MERS-CoV VLPs was restored in *NONO* KO + DPP4 cells compared to control *NONO* KO cells as measured by luciferase levels (Figure 3G). VLP entry in WT + DPP4 and *NONO* KO + DPP4 cells were equivalent and higher than in untransduced WT cells consistent with higher surface expression levels of DPP4. Transduction of DPP4 into WT cells also increased MERS-CoV titers in supernatants and recovered MERS-CoV titers in *NONO* KO cells to match the baseline titers recovered from WT cells (Figures 3H and S2D). However, despite equivalent surface expression of DPP4 following lentivirus transduction, MERS-CoV titers were 5-to 9-fold lower in *NONO* KO + DPP4 cells compared to WT + DPP4 cells. Expression of MERS-CoV N protein in the *NONO* KO + DPP4 cells was restored to WT levels by 24 hpi (Figure 3I). Additionally, while transcription of MERS-CoV genomic RNA was equivalent in DPP4-transduced WT and *NONO* KO cells over 72 hpi, accumulation of *M* subgenomic mRNA was reduced at this later timepoint in the absence of NONO (Figure 3J). We next investigated whether NONO directly interacts with MERS-CoV vRNA. Nuclear-cytoplasmic fractionation identified a significant amount of cytoplasmic-resident NONO protein in HAP1 cells that did not redistribute during MERS-CoV infection (Figure S2E). This cytoplasmic population of NONO did not appreciably bind MERS-CoV genomic vRNA or subgenomic mRNA species (2-to 3-fold enrichment) (Figure S2F). These findings suggest that the MERS-CoV replication defect in *NONO* KO cells is almost entirely due to a failure to appropriately transcribe and express sufficient DPP4 to facilitate optimal viral attachment and entry. However, a small defect remains apparent in peak virus titers in multi-cycle replication curves for *NONO* KO + DPP4 cells, suggesting a second role for NONO in MERS-CoV replication.

### NONO does not influence splicing or stability of DPP4 mRNA

NONO participates in post-transcriptional regulation of cellular gene expression by binding pre-mRNAs to enhance splicing. Indeed, defects in expression of several host genes required for mRNA splicing were observed in our RNA-seq datasets (Figure S1B). To further understand whether NONO impacts post-transcriptional regulation of *DPP4* mRNA, we performed replicate multivariate analysis of transcript splicing (rMATS) on our RNA-seq data, which quantifies alternative splicing events and identifies differentially regulated exons across samples. Our analysis identified skipped exons as the most significant alternative splicing events in *NONO* KO cells confirming our previous study that a dysregulation of splicing does occur (Cite) (Figure 4A). However, despite this general defect in host gene splicing in *NONO* KO cells, no significant changes in alternative splicing events were observed in *DPP4* mRNA isoforms (Figure 4B), and no accumulation of *DPP4* pre-mRNA was detected (Figure 4C and 4D). NONO has been previously described to regulate STAT3 expression through stabilizing interactions with the *STAT3* mRNA to prevent mRNA decay ^6^. To determine if NONO impacts *DPP4* mRNA half-life, we employed an actinomycin D RNA synthesis inhibition assay to track mRNA decay rates. An initial RNA immunoprecipitation confirmed endogenous NONO protein can bind *DPP4* mRNA (Figures S3A and S3B). However, actinomycin D treatment did not show a difference in DPP4 mRNA decay rates between WT and *NONO* KO cells over time (Figure 4E). Thus, the function of NONO in DPP4 expression is independent of its general cellular roles in mRNA splicing and stabilization.

**Figure 4.**
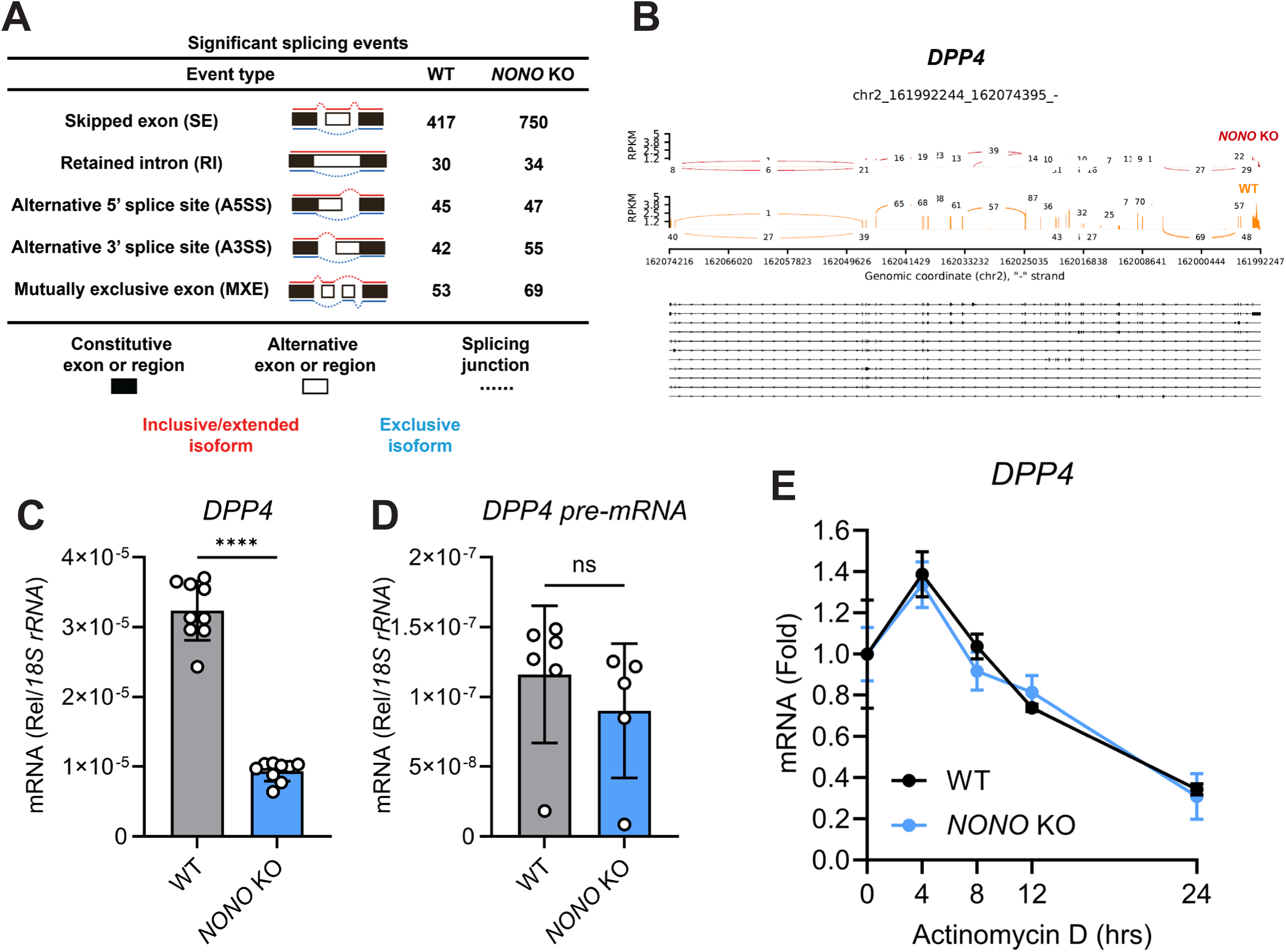
NONO does not influence splicing or stability of DPP4 mRNA: (A) rMATS alternative splicing events between WT and *NONO* KO HAP1 cells. (B) Sashimi plots for *DPP4* from WT or *NONO* KO HAP1 cells. (C-D) Mature and pre-mRNA expression for *DPP4* from WT or *NONO* KO HAP1 cells (qRT-PCR). (E) *DPP4* mRNA expression following actinomycin D treatment from WT or *NONO* KO HAP1 cells (qRT-PCR). Data are expressed as means (n=3) ± SD, ****p < 0.0001 (Welch’s t-test) and are representative of 2-3 independent experiments.

### NONO couples chromatin remodeling to expression of the MERS-CoV entry receptor

To determine if NONO alters the transcriptional state of the *DPP4* locus, we examined genome-wide differences in chromatin accessibility between WT and *NONO* KO cells using ATAC-seq. Chromatin accessibility within 3.0 Kb of the transcriptional start site was nearly identical between WT and *NONO* KO cells confirming previous studies that NONO itself is not a major chromatin remodeling component (Cite) (Figures 5A and 5B). Chromatin sites corresponding to the DPP4 locus were significantly less available in *NONO* KO cells matching our RNA-seq analysis (Figure 5C). Surprisingly, an Integrative Genomics Viewer track depicting the genomic region containing *DPP4* was unremarkable between WT and *NONO* KO samples except for a peak with reduced chromatin accessibility in the *NONO* KO near a non-coding region between exons 23 and 24 mapping to an enhancer region at the far 3’ end (Figures 5D). HOMER *de novo* motif discovery was used to identify transcription factor binding sites whose chromatin accessibility was impacted by NONO deletion.

**Figure 5.**
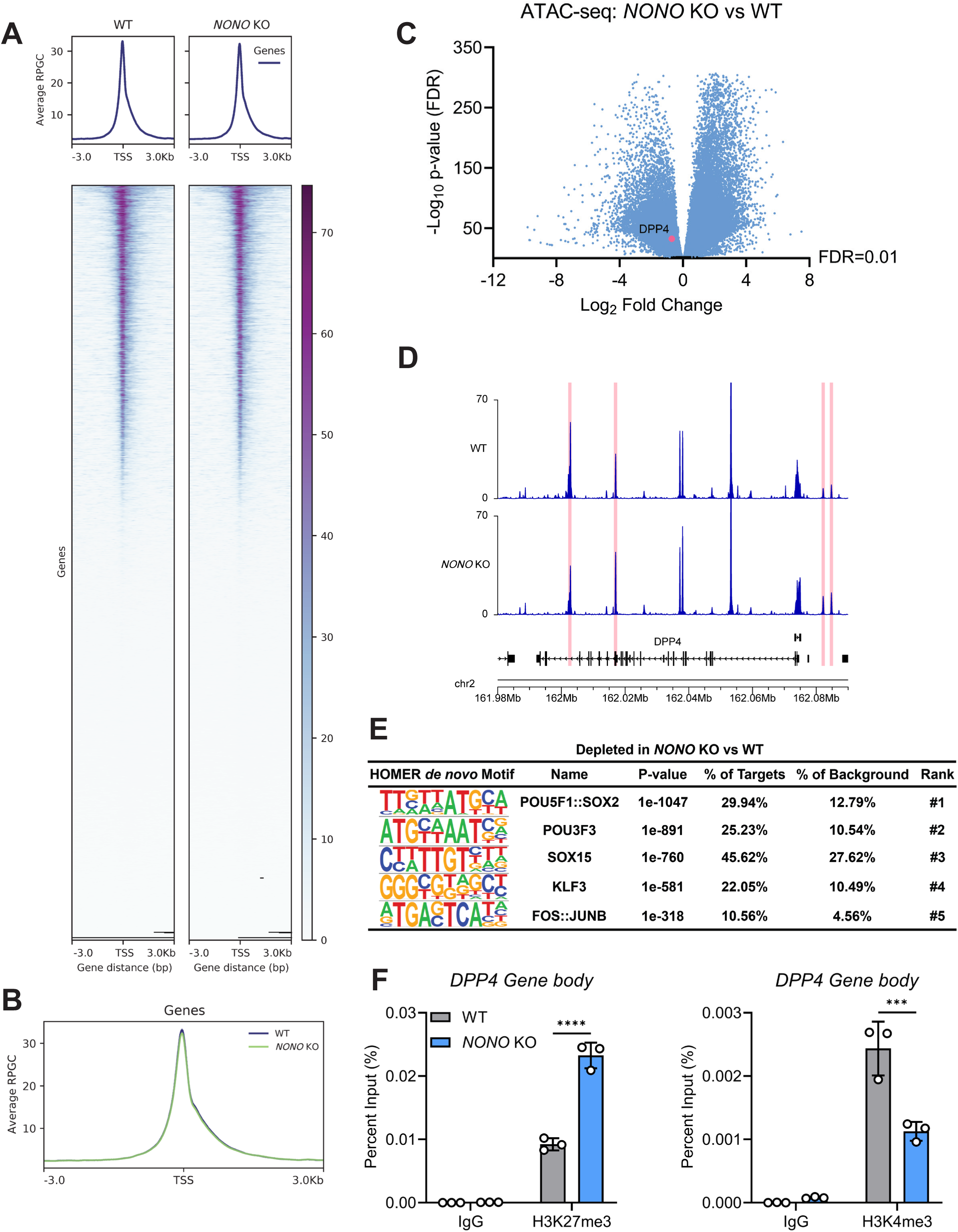
NONO couples chromatin remodeling to expression of the MERS-CoV entry receptor: (A-D) ATAC-seq of WT or *NONO* KO HAP1 cells (n=4 per group). (A) ATAC-seq signal for genome-wide chromatin accessibility (signal shown ± 3 Kb from TSS). (B) Average ATAC-seq profile plots for chromatin accessibility (signal shown ± 3 Kb from TSS). (C) ATAC-seq volcano plot for chromatin peaks of genes significantly less accessible in *NONO* KO compared to WT samples. (D) Genome browser view of open chromatin peaks for *DPP4* with highlighted portions indicating regions with significantly less available chromatin in the *NONO* KO compared to the WT. (E) HOMER *de novo* motif analysis for depleted motifs in *NONO* KO compared to WT samples. (F) ChIP-qPCR showing either H3K4me3 or H3K27me3 histone modifications at the *DPP4* gene body from WT or *NONO* KO HAP1 cells. Data are expressed as means (n=3) ± SD, ***p < 0.001, ****p < 0.0001 (two-way ANOVA with Sidak’s multiple comparisons) and are representative of 2-3 independent experiments.

NONO KO cells showed depleted accessibility for motifs corresponding to components of the AP-1 complex (FOS and JUNB) (Figure 5E). DPP4 contains neither a TATAA nor a CCAAT box as a promoter but has a C- and G-rich region containing several consensus binding sites for transcriptional factors including components of AP-1. To verify that reduced chromatin accessibility was responsible for the DPP4 transcription loss in *NONO* KO cells, we performed chromatin immunoprecipitation (ChIP) targeting histone modifications for either an open, transcriptionally active chromatin state (H3K4me3) or a closed, transcriptionally repressed form (H3K27me3) followed by qRT-PCR. H3K27me3 levels on the *DPP4* gene body were significantly enhanced while H3K4me3 markers were reduced in *NONO* KO cells (Figure 5F). These histone changes in the *NONO* KO cells were specific for DPP4 and were not observed for unrelated control genes including the actively transcribed RPL30 gene and transcriptionally silent α satellite repeat element (Figure S3C). Taken together, our data demonstrates that NONO is essential for proper chromatin architecture and gene expression for the MERS-CoV entry receptor DPP4.

## Discussion

The paraspeckle protein NONO is a multifunctional regulator of diverse biological activities including gene regulation, cell proliferation, apoptosis, migration, and DNA damage repair ^7^. In this study, we report a new role for NONO as a host factor required for the expression of the MERS-CoV entry receptor DPP4. Deletion of NONO severely limited MERS-CoV replication in an IFN-independent manner. MERS-CoV was unable to initiate vRNA replication and transcription in the absence of NONO which was due to impaired entry. NONO influences DPP4 transcription not by splicing or stability of the DPP4 mRNA, but by altering the balance of transcriptionally active and inactive histone modifications at the DPP4 locus. This results in reduced protein and cell surface expression of DPP4 leading to diminished MERS-CoV susceptibility in *NONO* KO cells.

Defining the factors that influence cell-surface receptors expression may prove beneficial for understanding viral host range, tissue and cell type susceptibility, and disease severity ^8^. Several risk factors associated with severe respiratory virus infections (increased age, smoking, and chronic obstructive pulmonary disease) also correlate with heightened expression of ACE2 and DPP4 ^9–13^. Although the molecular mechanisms governing these phenomena remain undefined, heightened entry receptor expression provides a biological rationale for the increased infection susceptibility in patients with these pre-existing conditions. Expression of viral entry receptors is a key determinant for pathogenesis. Recent studies have revealed that the SWItch/Sucrose Non-Fermentable (mSWI/SNF) chromatin remodeling complex is essential for chromatin accessibility at the ACE2 locus, ACE2 expression, and SARS-CoV-2 susceptibility ^8^. The distribution of DPP4 in host cells significantly impacts the transmission potential of MERS-CoV as well as the severity of the infection. MERS-CoV is not efficiently transmitted between humans, but it is highly prevalent in the dromedary camel reservoir host with seroprevalence levels greater than 71% in 4 separate studies of dromedary populations epidemiologically linked with human MERS-CoV infection ^14^. This inefficient transmission is largely due to an absence of DPP4 in the human upper respiratory tract epithelium while dromedary camels retain high expression levels of the receptor. Despite this, MERS-CoV superspreader events have occurred through human-to-human transmission in healthcare settings, raising concern for the potential of sustained chains of viral transmission in humans ^15,16^. The human lower respiratory tract has relatively higher expression levels of DPP4 with primary human airway epithelia from 74 individual donors revealing significant inter-individual variation in DPP4 expression ^17^. This spectrum in DPP4 abundance correlated with MERS-CoV infection outcomes where more DPP4 allowed for early, robust virus replication and increased infection severity.

Surprisingly, early efforts to generate mouse models for MERS-CoV revealed homozygous transgenic mice expressing human DPP4 were unexpectedly more resistant than heterozygous mice to MERS-CoV infection. This resistance was attributed to an increase in soluble human DPP4 present in the serum which retained virus neutralizing activity ^18^. The implication that viral entry receptor expression can be as beneficial as it is harmful has prompted new investigations into engineered ACE2 and DPP4 decoy receptors as a robust therapeutic for SARS-CoV-2 and MERS-CoV infection respectively ^19,20^. In addition to entry receptor saturation, it may also be beneficial to explore inhibition of key transcriptional components required for entry receptor expression as a strategy to limit viral replication. Electrophilic compound screening recently identified (R)-SKBG-1 as a covalent NONO-specific ligand that suppress cancer-relevant gene transcription and cancer cell proliferation ^21^. (R)-SKBG-1 was determined to suppress protumorigenic transcriptional networks by accumulating NONO in nuclear foci and stabilizing NONO-RNA interactions thereby trapping mRNA processing. Co-opting of host RBPs and transcription factors by small molecules may provide a method to selectively repress expression of disease-relevant proteins.

Several host RBP-SARS-CoV-2 vRNA interactome screens have now been published with NONO identified in a consensus set of 58 RBPs between the different studies ^1–3^. Interestingly, the related DBHS paraspeckle protein SFPQ was also enriched across these three screens possibly due to the requirement for homo-or heterodimerization between the three DBHS members for proper cellular function. SFPQ is involved in modulating viral transcription and replication for influenza A virus by polyadenylating viral mRNAs ^22^ and for repressing expression of Epstein-Barr virus viral lytic genes by promoting histone H1 occupancy at the viral genome ^23^. These studies, combined with the identification of both NONO and SFPQ across three different RBP-SARS-CoV-2 vRNA-interactome screens, implicate DBHS proteins as host factors that are more commonly involved in vRNA regulation than previously known. NONO and SFPQ are predominantly localized in the nucleus while CoV replication occurs in the cytoplasm, although SFPQ nuclear-cytoplasmic translocation has been reported during encephalomyocarditis virus infection to support viral replication ^24^. Our data identifies a sizable cytoplasmic portion of NONO present in HAP1 cells under basal conditions and that MERS-CoV infection does not alter NONO distribution between the nuclear and cytoplasmic compartments. Despite this high cytoplasmic presence, NONO modestly bound MERS-CoV vRNA suggesting NONO may not play a major role in MERS-CoV genome replication. We recently reported a critical role for NONO in regulating the antiviral innate immune response by promoting chromatin accessibility, recruitment of RNA polymerase II to *IFNB1* and ISG promoters, and transcription of antiviral genes ^4^. While we do not observe effects on antiviral gene transcription due to the viral antagonism on those pathways, it is still possible that usurping of NONO by MERS-CoV for genome replication may reduce innate immune activation under certain circumstances.

Whether NONO has a direct role in SARS-CoV-2 or MERS-CoV genome replication remains to be determined and future studies will be necessary to delineate these mechanisms. *NONO* KO + DPP4 cells displayed 5-to 9-fold lower MERS-CoV titers compared to WT + DPP4 cells indicating an additional replication impairment beyond DPP4 levels persists. A possible explanation for this gap in titers is the requirement for host lipids for CoV replication. NONO has been described maintaining sterol regulatory element-binding protein (SREBP)-Regulated cholesterol biosynthesis ^25^ and reprogramming of the SREBP-regulated lipid biosynthesis pathways is indispensable for satisfying MERS-CoV and SARS-CoV-2 lipid demands during replication ^26–30^. The GSEA pathways regulation of cholesterol biosynthesis by SREBP and activation of gene expression by SREBF were both significantly down regulated in *NONO* KO cells and are in agreement with prior published RNA-seq datasets highlighting the importance of NONO for expression of SREBP targets ^25^.

In summary, our study identifies NONO as a MERS-CoV host dependency factor by regulating the expression of the MERS-CoV entry receptor DPP4. Our findings provide new evidence on the extent RBPs like NONO play in infection and disease. DBHS proteins like NONO can greatly influence gene expression through chromatin regulation, pre-mRNA splicing, and paraspeckle formation making them appealing targets for pharmacological intervention. By developing a clearer understanding of the host factors and mechanisms that regulate expression of viral entry receptors, we may inform the development of better countermeasures and therapeutics.

## Acknowledgments

This research was supported by the Intramural Research Program of the National Institutes of Health (NIH). The contributions of the NIH authors are considered Works of the United States Government. The findings and conclusions presented in this paper are those of the authors and do not necessarily reflect the views of the NIH or the U.S. Department of Health and Human Services.

## Author contributions

A.H. performed all aspects of this study. M.J. and S.D.S performed experiments. L.M.M, A.B.C., L.T., O.S., and K.L.M provided critical reagents and technical advice. S.Y., K.R., T.E.M., P.A.B., J.B.L., and C.M. performed analysis of sequencing datasets. A.H. and S.M.B. organized the study and prepared the manuscript. All authors have read and agreed to the published version of the manuscript.

## RESOURCE AVAILABILITY

### Lead contact

Further information and requests for resources and reagents should be directed to and will be fulfilled by the lead contacts, Adam Hage and Sonja M. Best.

## Materials availability

Resources and reagents generated in this study are available upon request from the lead contacts.

## Data and code availability

Transcriptomic and epigenomic data generated during this study have been deposited to the NCBI Gene Expression Omnibus (GEO) database under GEO: GSE348212 (RNA-seq) and GEO: GSE347963 (ATAC-seq). All datasets are publicly available as of the date of publication.

This paper does not report original code.

Any additional information required to reanalyze the data reported in this study is available from the lead contact upon request.

## EXPERIMENTAL MODEL AND SUBJECT DETAILS

### Cell lines

HAP1 cell lines WT (C631), *NONO* KO (HZGHC007548c001), and *DPP4* KO (HZGHC003881c011) were purchased from Horizon Discovery and maintained in Iscove’s Modified Dulbecco’s Medium (IMDM) (Gibco) supplemented with 10% (v/v) fetal bovine serum (FBS) (Gibco) and 1% (v/v) penicillin-streptomycin (Gibco). Vero (CCL-81) and HEK293T (CRL-3216) purchased from ATCC and A549s expressing human ACE2 and TMPRSS2 (A549 hACE2/hTMPRSS2) kindly provided by Jonathan W. Yewdell (NIH/NIAID) were maintained in Dulbecco’s Modified Eagle’s Medium (DMEM) (Gibco) supplemented with 10% (v/v) fetal bovine serum (FBS) (Gibco) and 1% (v/v) penicillin-streptomycin (Gibco). Cells used for transfections were plated in media supplemented with 10% (v/v) FBS lacking 1% (v/v) penicillin-streptomycin.

### Viruses

Viruses used in this study were handled under biosafety level 3 (BSL-3) conditions at the Rocky Mountain Laboratories Integrated Research Facility in accordance with Division of Select Agents and Toxins (DSAT) regulations for study of select agents and Institutional Biosafety approvals. SARS-CoV-2 (USA-WA1/2020) was kindly provided by The World Reference Center of Emerging Viruses and Arboviruses (WRCEVA) (The University of Texas Medical Branch at Galveston). MERS-CoV (HCoV-EMC/2012) was obtained from BEI Resources.

## METHOD DETAILS

### Inhibitors

IFN-I signaling blockade was performed with Ruxolitinib or DMSO control diluted in IMDM to a concentration of 10 μg/mL (InvivoGen). Cells were pretreated with Ruxolitinib or DMSO for 1 hour before infection. Transcription inhibition and mRNA stability assays were performed with actinomycin D or DMSO control diluted in IMDM to a concentration of 1 μM (Cell Signaling Technology). Compounds were maintained at the indicated concentration in culture for the duration of the experiment.

### Immunoblot assay

Cell lysates were resolved on 10% Novex Tris-Glycine gels (Invitrogen) and transferred to polyvinylidene difluoride (PVDF) membranes using the Trans-Blot Turbo transfer system (Bio-Rad). Membranes were blocked with 5% (w/v) non-fat dry milk in TBST (TBS with 0.1% (v/v) Tween-20) for 1 hour, washed with TBST three times for 5 minutes each, and probed with the indicated primary antibody in 3% (w/v) BSA in TBST at 4°C overnight. Following overnight incubation, membranes were washed with TBST three times for 5 minutes each, probed with anti-rabbit or anti-mouse IgG (whole molecule)-peroxidase antibody produced in goat (Sigma) in 5% (w/v) non-fat dry milk in TBST for 1 hour at room temperature, and washed with TBST three times for 5 minutes each. Proteins were visualized by ECL (Pierce) or SuperSignal West Femto chemiluminescence reagents (Thermo Scientific) and detected using an iBright FL1500 Imaging System (Invitrogen). The primary antibodies and concentrations used are listed in the key resources table.

### Infections and plaque assays

HAP1 cells were seeded in 24-well plates (150,000 cells/well) and infected with virus diluted in IMDM (DMEM for A549 hACE2/hTMPRSS2 cells) at 37°C for 1 hour. Synchronized infections were performed at 4°C for 1 hour. Inoculations were removed and cells were washed once with DPBS, overlaid with media containing 2% (v/v) FBS, and incubated at 37°C. Supernatants were collected for plaque assay at the indicated time points. For plaque assays, confluent monolayers of Vero cells were inoculated with supernatants serially diluted in DMEM containing 2% (v/v) FBS and 1% (v/v) penicillin streptomycin and incubated at 37°C for 1 hour.

Inoculums were removed and replaced with MEM containing 1.5% (w/v) carboxymethylcellulose and 1% (v/v) penicillin streptomycin and incubated at 37°C for 3 days (4 days for SARS-CoV-2). Cells were fixed in 10% (w/v) formalin for 1 hour at room temperature and stained with 1% (w/v) crystal violet for 10 minutes at room temperature.

### Quantitative reverse transcription PCR (qRT-PCR)

Total RNA isolation and gDNA removal was attained using the RNeasy Plus Mini Kit with gDNA Eliminator columns (Qiagen). cDNA synthesis was achieved using SuperScript VILO Master Mix (Invitrogen). qRT-PCR was performed using SsoAdvanced Universal SYBR Green Supermix (Bio-Rad) in a 384-well QuantStudio 7 Pro Real-Time PCR System (Applied Biosystems). Gene expression was normalized to human 18S rRNA by the comparative CT method (ΔΔCT). The primer sequences used are listed in Table S1.

### Virus-like particle entry assay

Virus-like particles (VLPs) for MERS-CoV (Virongy Biosciences) consisted of an alphavirus vector RNA including a firefly luciferase reporter gene and the MERS-CoV spike, envelope, nucleocapsid, and membrane proteins. WT and *NONO* KO HAP1 cells were seeded in 96-well plates (25,000 cells/well) and infected with 15 μL of IMDM and 45 μL of either the MERS-CoV VLPs or IMDM control. After 6 hours, cells were washed with DPBS, lysed in 20 μL of passive lysis buffer, and incubated at room temperature on a platform rocker for 15 minutes. Luciferase activity was measured using the Dual-Luciferase Reporter Assay System on a GloMax Explorer (Promega) according to the manufacturer’s instructions.

### Chromatin immunoprecipitation (ChIP) and ChIP-qPCR

ChIP was performed using the SimpleChIP Plus Sonication Chromatin IP Kit (Cell Signaling Technology) according to the manufacturer’s instructions. 2 μg of ChIP-grade antibodies were used for each immunoprecipitation. ChIP-qPCR was performed using SimpleChIP Universal qPCR Master Mix (Cell Signaling Technology) in a 384-well QuantStudio 7 Pro Real-Time PCR System (Applied Biosystems). Immunoprecipitation efficiency was calculated using the percent input method (2% x 2^(CT^ ^2%^ ^Input^ ^Sample-CT^ ^IP^ ^Sample)^). The primer sequences used are listed in Table S1.

### RNA immunoprecipitation (RIP)

HAP1 cells were seeded in 10 cm dishes (3 x 10^6^ cells/sample) and treated as indicated. Cells were harvested in RIP lysis buffer (25mM Tris-HCl, pH 7.4, 150mM KCl, 5mM EDTA, 0.5mM DTT, 0.5% (v/v) IGEPAL CA-630, 100 U/mL SUPERase·In RNase Inhibitor, and protease inhibitor cocktail (Roche). Cell lysates were clarified by centrifugation at 21,000 x RCF for 20 min at 4°C and 10% of the clarified lysate was added to 2X Laemmli buffer containing 2-Mercaptoethanol, heated for 10 min at 95°C, and RLT Plus RNA extraction buffer as whole-cell lysate (WCL) protein and RNA inputs. The remaining lysate was subjected to immunoprecipitation with 2 μg of primary antibody overnight at 4°C followed by incubation with protein A/G agarose beads (Pierce) for 2 hours at 4°C on a rotating platform. Beads were washed seven times with RIP buffer and 10% of the bead slurry was collected for protein inputs. The remaining beads were resuspended in RLT Plus RNA extraction buffer to isolate bound RNAs.

### Lentivirus production and generation of DPP4 stable cells

To generate lentiviruses, HEK293T cells were seeded in 10 cm dishes (1.5 x 10^6^ cells/dish) and transfected with 2.5 μg of the DPP4 lentiviral vector, 2 μg of psPAX2, 0.8 μg of pMD2.G, and 15.9 μL of Mirus LT1 transfection reagent (Mirus) prepared in 250 μL of Opti-MEM (Gibco) according to the manufactures recommendation. Forty-eight hours post-transfection, lentivirus supernatants were collected, centrifuged at 1000 x RCF for 5 minutes to pellet cell debris, and aliquots stored at −80°C. Collected lentiviruses were reverse transduced into *NONO* KO HAP1 cells seeded in 6-well plates (50,000 cells/well) for 48 hours in the presence of 10 μg/mL of polybrene (Sigma). Transduction media was replaced with antibiotic selection media containing 5 μg/mL blasticidin (InvivoGen). Cells remained under selection until all non-transduced cells were dead. Successful DPP4 transduction and cell surface expression was confirmed by flow cytometry, immunoblot, and qRT-PCR.

### Flow cytometry

WT and *NONO* KO HAP1 cells (1 x 10^6^ cells/sample) were centrifuged at 500 x RCF for 5 minutes to pellet cells and remove supernatant. Cells were washed with DPBS and resuspend in 100 μL of live/dead stain (Invitrogen) diluted 1:500 in DPBS for 30 minutes at room temperature protected from light. Live/dead stain was quenched with 500 μL of wash buffer (2% (v/v) FBS in DPBS) and washed again with 500 μL of wash buffer. Cells were resuspended in 100 μL of DPP4 (Invitrogen) or IgG1 (BD Biosciences) primary antibody diluted 1:50 in wash buffer and incubated at room temperature for 1 hour. Cells were washed in 500 μL of wash buffer and resuspended in 100 μL of fluorochrome-conjugated secondary antibody (Invitrogen) diluted 1:100 in wash buffer. Cells were incubated for 30 minutes at room temperature and washed in 500 μL of wash buffer before fixation in 200 μL of 2% (v/v) paraformaldehyde for 30 minutes at room temperature. Cells were washed in 500 μL of DPBS and resuspended in 200 μL of wash buffer for analysis on a BD FACSymphony A5 flow cytometer.

### Cellular fractionation

Nuclear and cytoplasmic compartments were separated using the NE-PER nuclear and cytoplasmic extraction kit (Thermo Scientific) according to the manufacturer’s instructions.

### NONO siRNA knockdown

Transient knockdown of endogenous NONO in A549 hACE2/hTMPRSS2 cells, seeded in 24-well plates (30,000 cells per well), was achieved by transfection of ON-TARGETplus Non-targeting Control Pool (D-001810-10-05 Dharmacon) or SMARTpool: ON-TARGETplus NONO siRNA (L-007756-01-0005 Dharmacon) for a final concentration of 20 nM siRNA. Delivery of siRNA was achieved with Lipofectamine RNAiMAX (Invitrogen) according to the manufacture’s guidelines.

### RNA-seq data processing

Total RNA isolation and gDNA removal was attained using the RNeasy Plus Mini Kit with gDNA Eliminator columns (Qiagen) according to the manufacturer’s instructions. RNA-seq libraries were prepared using the NEBNext Ultra II DNA Library Prep Kit according to the manufacturer’s instructions. RNA-seq libraries were sequenced on a NextSeq 2000 P2 using Illumina mRNA Ligation Kit and paired-end sequencing. The samples have 158 to 188 million pass filter reads with more than 93% of bases above the quality score of Q30. Reads were trimmed for adapters and low-quality bases using Cutadapt version 4.4 ^31^ and aligned to the human reference genome (hg38) and Gencode 30 annotation using STAR in single-pass mode ^32^. The average mapping rate across all samples was 96%, with unique alignment above 89% and 2.69-4.37% unmapped reads. Mapping statistics were calculated using Picard software. The samples contained 0.72% ribosomal bases; coding bases ranged from 63-64%, UTR bases from 32-32%, and mRNA bases from 96-96% across all samples. Library complexity was measured in terms of unique fragments in the mapped reads using Picard’s MarkDuplicates utility, with samples containing 56-58% non-duplicate reads. Gene expression quantification was performed for all samples using RSEM ^33^. Differential gene expression analysis was performed using limma-voom ^34^ as implemented in eVITTA, a web-based visualization and inference toolbox for transcriptome analysis ^35^. Alternative splicing events were quantified using rMATS (v4.3.0) ^36^, which detects and statistically evaluates five categories of alternative splicing events - skipped exons (SE), alternative 5′ splice sites (A5SS), alternative 3′ splice sites (A3SS), mutually exclusive exons (MXE), and retained introns (RI) - from replicate RNA-seq data. Differential splicing events from all five categories were filtered by FDR < 0.05 and |ΔPSI| ≥ 0.10, collapsed to the gene level, and ranked by ascending FDR and descending |ΔPSI| to select the top 10 genes for downstream analysis.

### ATAC-seq data processing

ATAC-seq sample and library preparation was performed on 100,000 fresh cells using the ATAC-seq Kit (Active Motif) according to the manufacturer’s instructions. ATAC-seq libraries were sequenced on a NovaSeq X Plus 1.5B flow cell using paired-end sequencing. The samples yielded 556 to 811 million pass-filter reads, with more than 92% of bases above a quality score of Q30. ATAC-seq reads were processed using the chrom-seek (2.0.0) pipeline. In brief, reads were trimmed with Cutadapt version 4.4 ^31^ and then aligned the human GRCh38.p12 genome using BWA version 0.7.17 ^37^. All reads aligning to the Encode hg38 v2 blacklist regions ^38^ were identified and removed with Picard SamToFastq. Reads with a mapQ score less than 6 were removed with SAMtools version 1.17 ^39^ and PCR duplicates were removed with Picard MarkDuplicates. Data was converted into bigwigs for viewing and normalized by reads per genomic content (RPGC) using deepTools version 3.5.5 ^40^. Averaged bigwigs as well as TSS-associated heatmaps and profile plots were also created using deepTools and ggplot2 version 4.0.3. Peaks were called using Genrich version 0.6. Consensus peaks across all conditions and associated PCA based upon peak intensities was calculated using DiffBind v2 ^41^.

Differential peaks were called using DiffBind v2 and its Deseq2 ^42^ differential caller with default parameters. Peaks were then annotated to nearest TSS using UROPA version 4.0.2 ^43^ and Gencode Release 28. Motif analysis was performed on differential peaks using HOMER v 4.11.1 ^44^ with default parameters. Volcano plot was created using GraphPad Prism 10. Example region plots were created using karyoploteR version 1.36.0.

## QUANTIFICATION AND STATISTICAL ANALYSIS

All data were presented as means ± SD and analyzed using GraphPad PRISM software (version 10.5.0 GraphPad Software). Welch’s t-test or two-way ANOVA with Sidak’s or Tukey’s multiple comparisons were used. *p < 0.05; **p < 0.01; ***p < 0.001; ****p < 0.0001.

## Key resources table

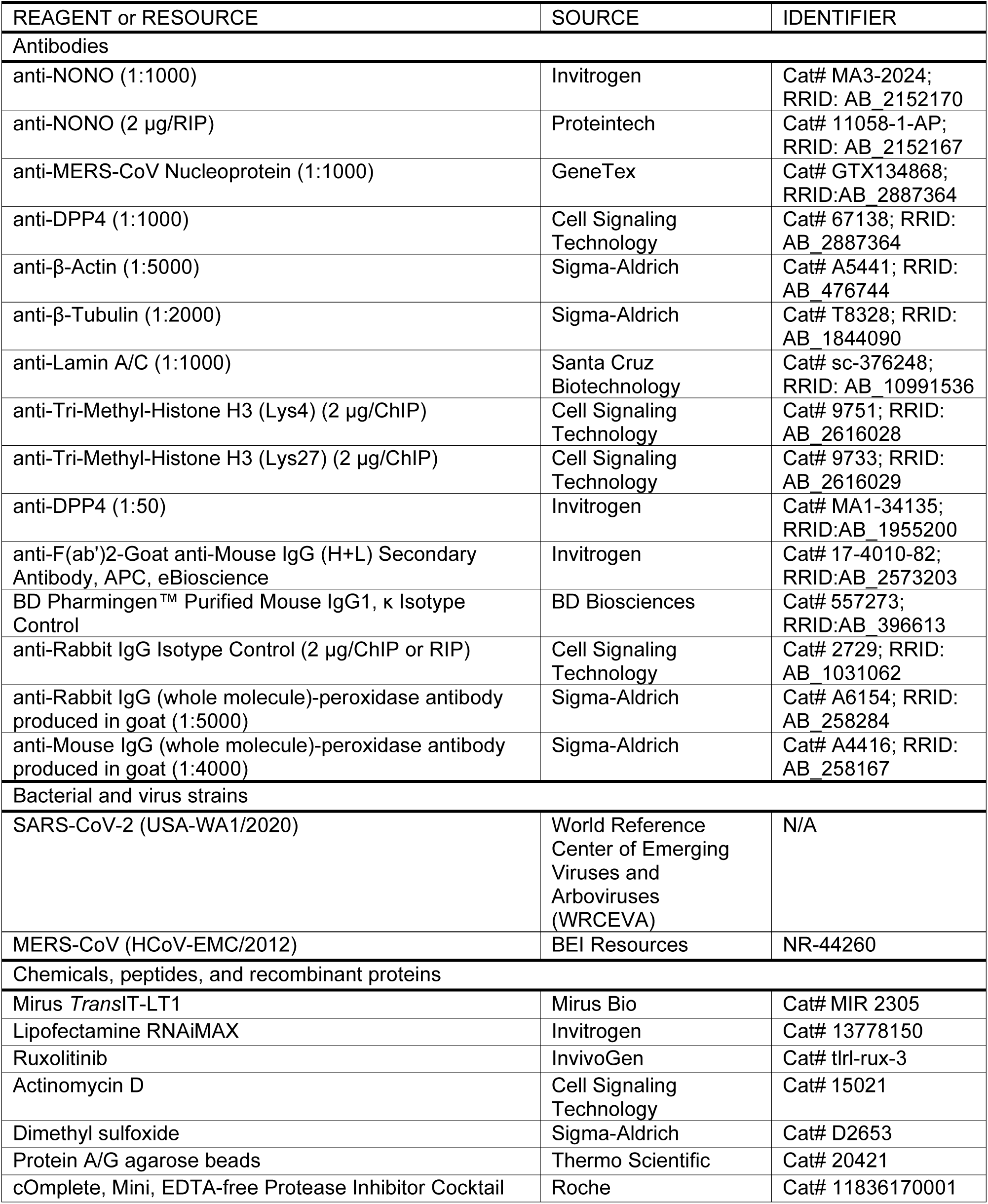
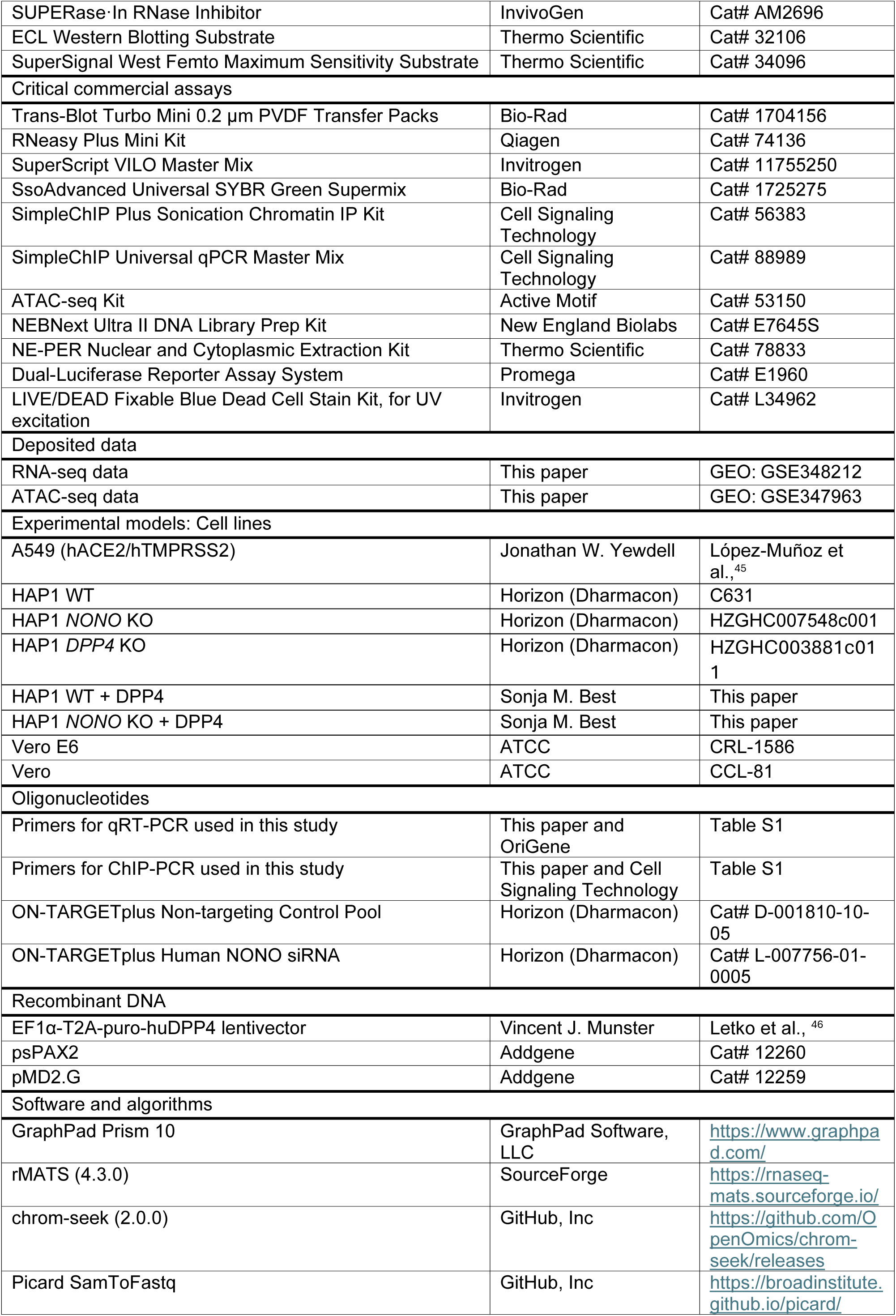
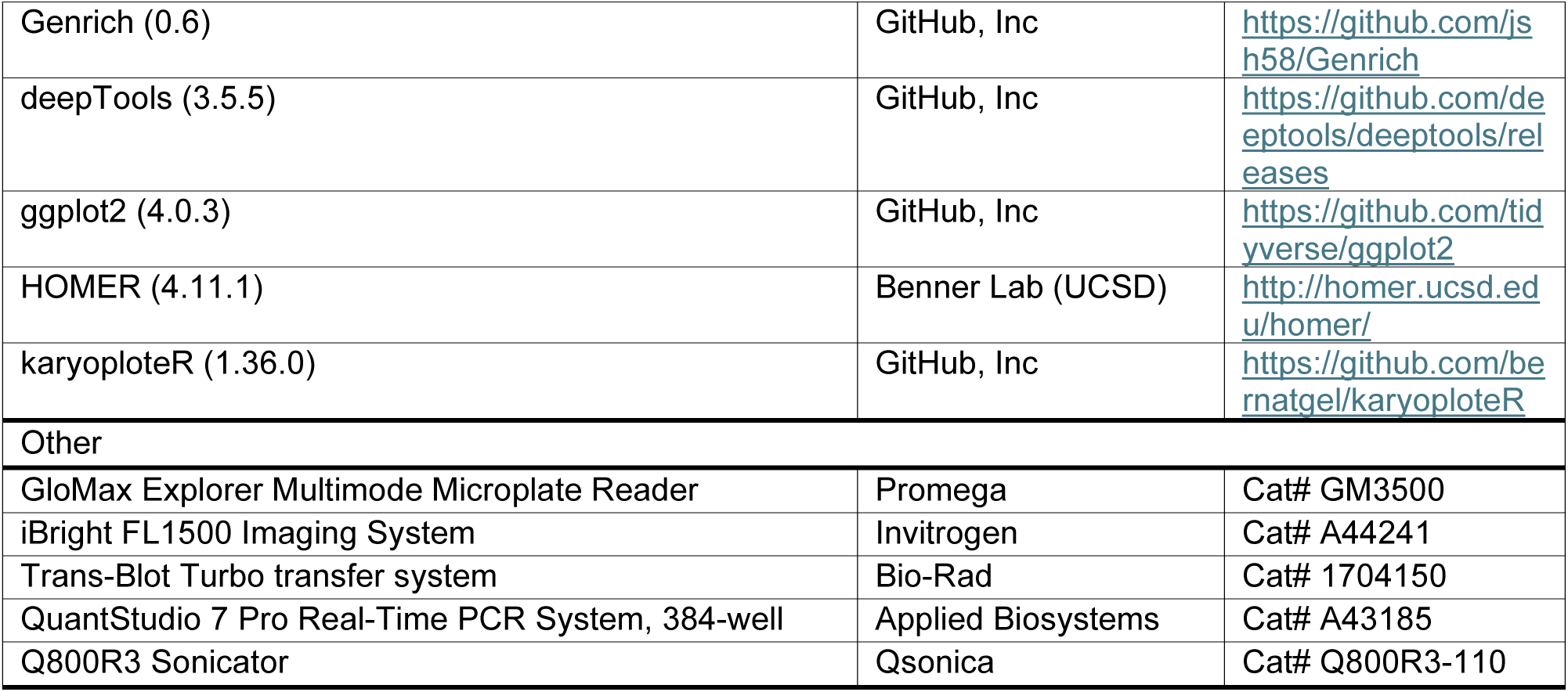

---

**Table S1:**
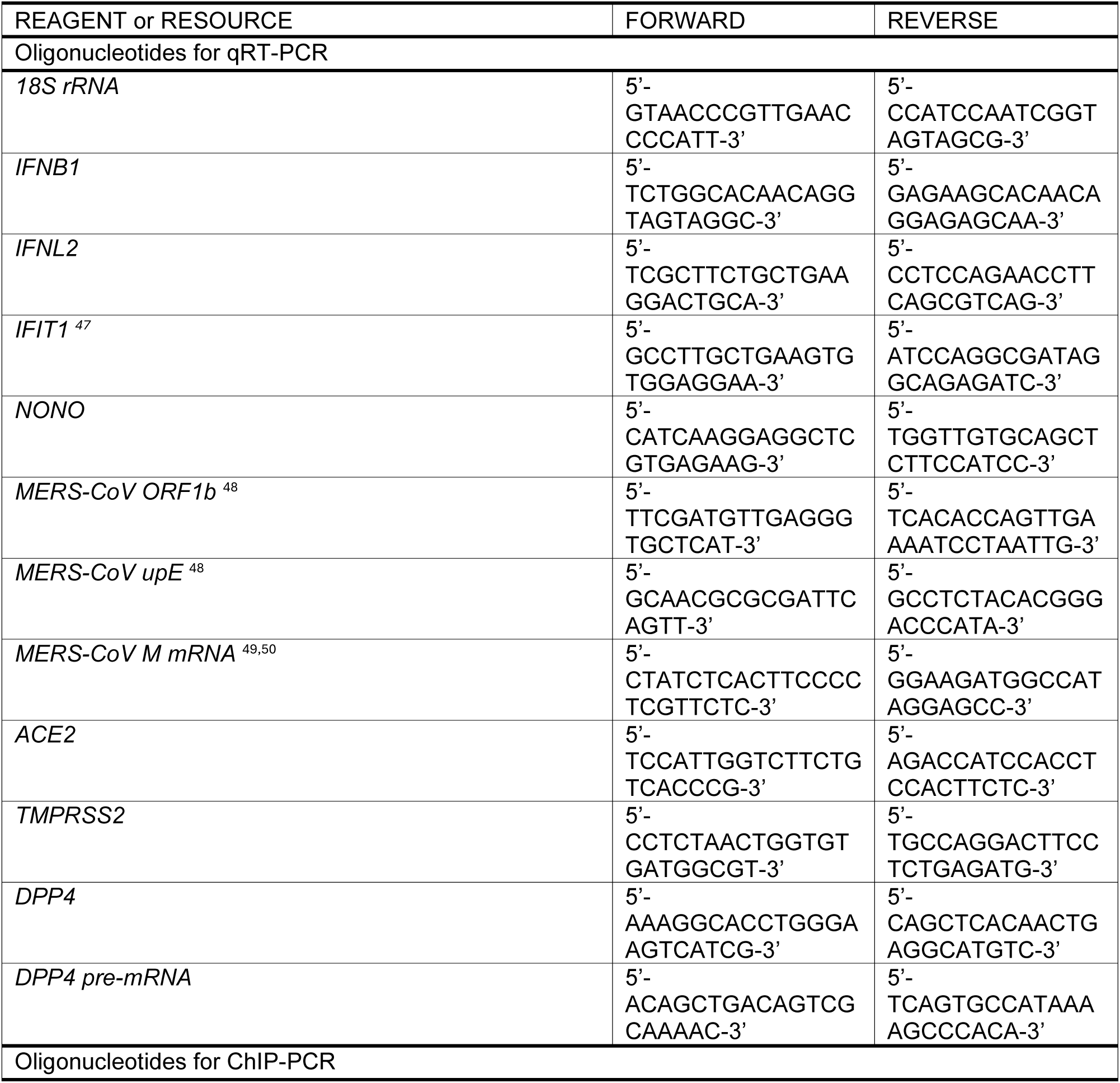
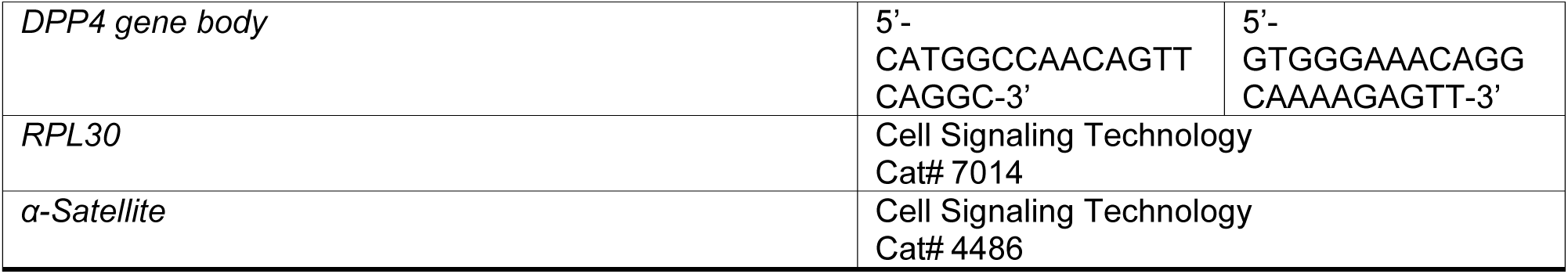
Primer sequences.

**Figure S1:**
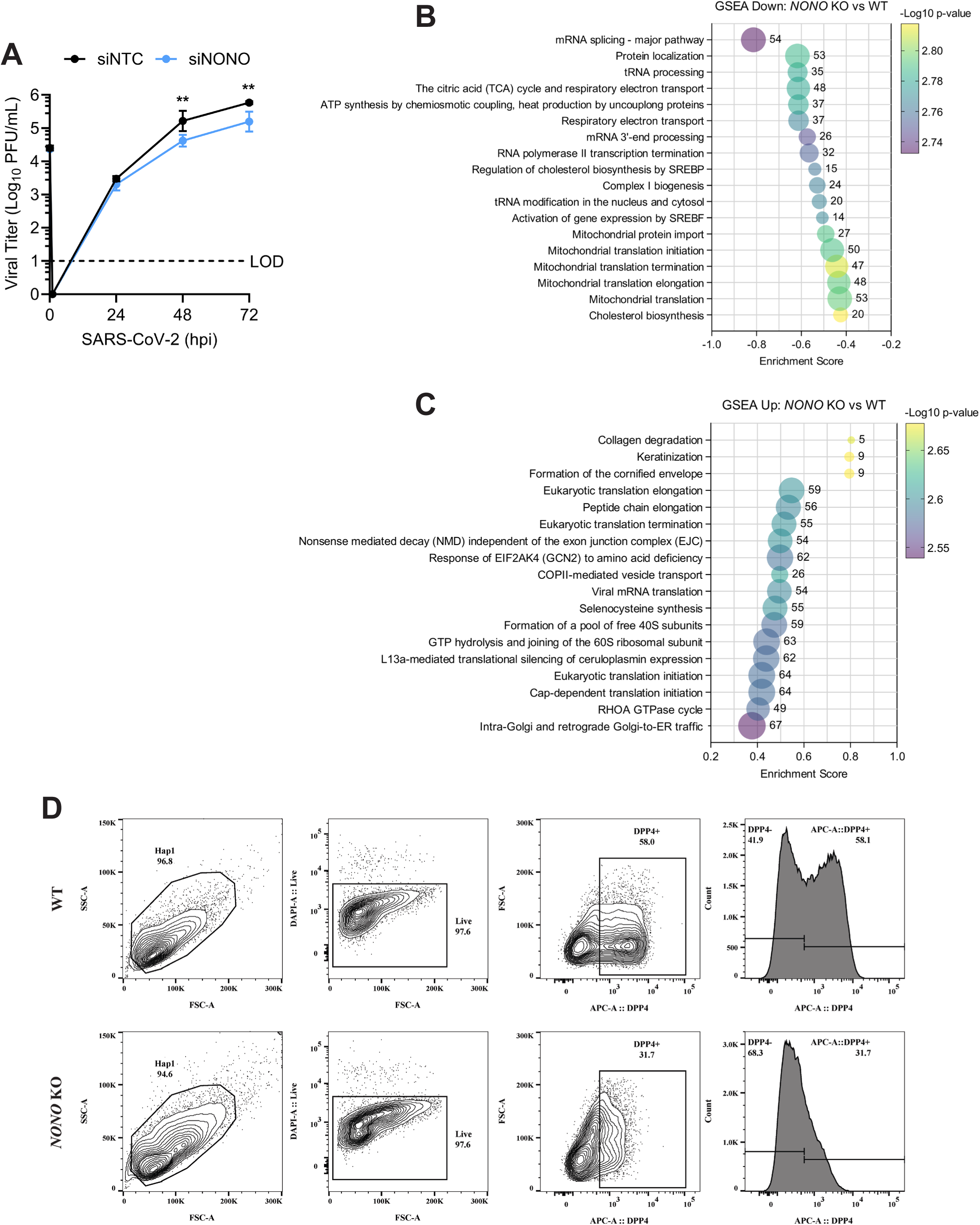
NONO is an essential host factor for MERS-CoV replication: (A) Infectious viral titers from hACE2/hTMPRSS2 expressing A549 cells treated with siNTC or siNONO infected with SARS-CoV-2 (MOI=0.01) (plaque assay). (B-C) GSEA terms and pathways of down and upregulated transcripts from WT or *NONO* KO HAP1 samples. Numbers adjacent to bubbles represent gene counts. (D) Gating strategy to determine DPP4 surface expression from WT or *NONO* KO HAP1 cells (flow cytometry). Data are expressed as means (n=3) ± SD, **p < 0.01, (two-way ANOVA with Sidak’s multiple comparisons) and are representative of 2-3 independent experiments.

**Figure S2:**
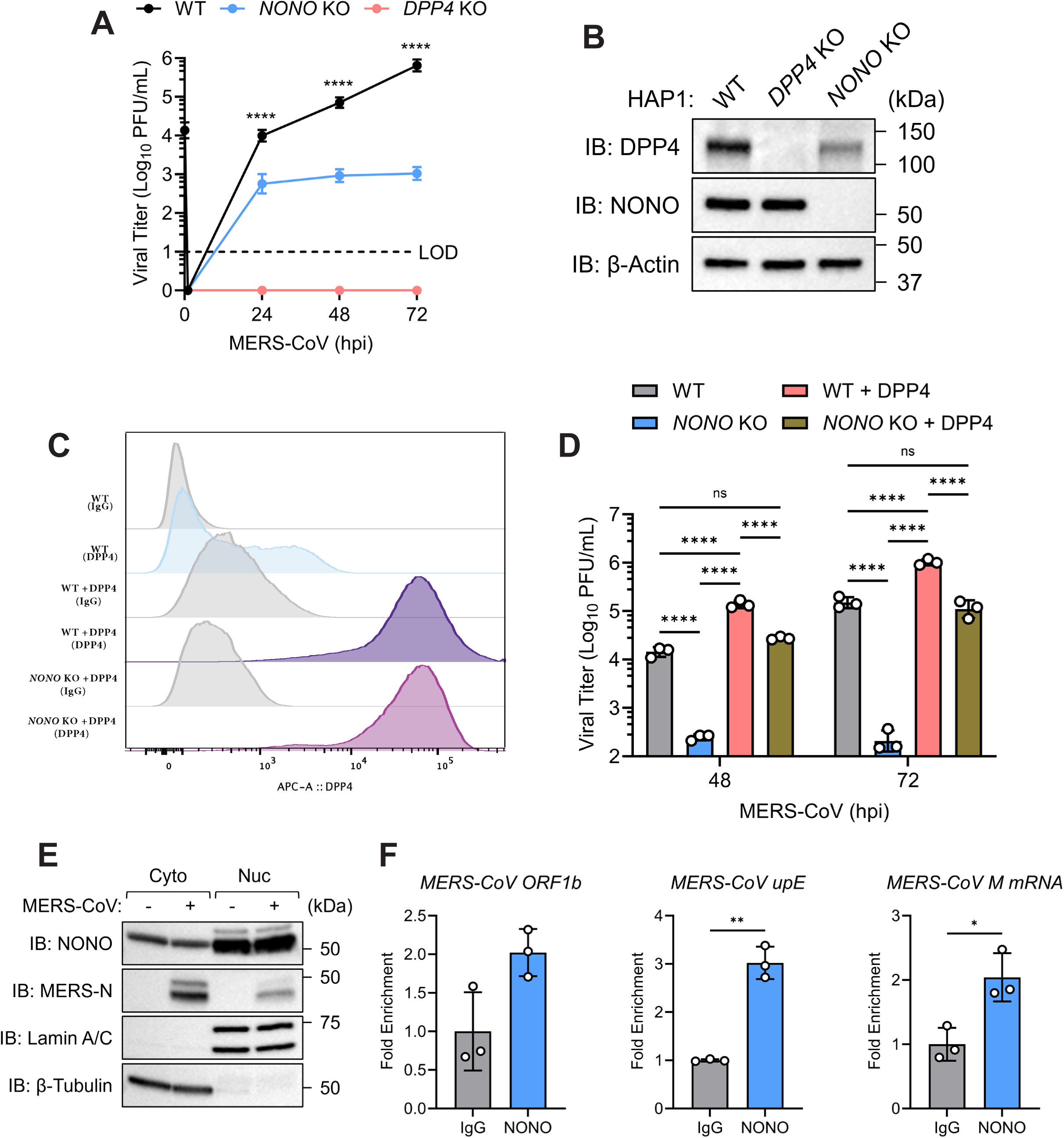
NONO deletion disrupts DPP4 transcription and MERS-CoV entry: (A) Infectious viral titers from WT, *NONO* KO, and *DPP4* KO HAP1 cells infected with MERS-CoV (MOI=0.01) (plaque assay). (B) Protein levels for DPP4 and NONO from WT, *NONO* KO, and *DPP4* KO HAP1 cells (immunoblot). (C) DPP4 surface expression from DPP4 transduced or untransduced WT or *NONO* KO HAP1 cells (flow cytometry). (D) WT or *NONO* KO HAP1 cells infected with MERS-CoV (MOI=0.01) zoom in of 48-72 hpi infectious viral titers from (Figure 3H) (plaque assay). (E-F) WT HAP1 cells infected with MERS-CoV (MOI=0.01) for 24 hours. (E) Subcellular fractionation (immunoblot). (F) Expression of NONO-bound MERS-CoV vRNA from endogenous NONO RNA immunoprecipitation (qRT-PCR). Data are expressed as means (n=3) ± SD, *p < 0.05, **p < 0.01, ****p < 0.0001 (two-way ANOVA with Dunnett’s or Tukey’s multiple comparisons or Welch’s t-test) and are representative of 2-3 independent experiments.

**Figure S3:**
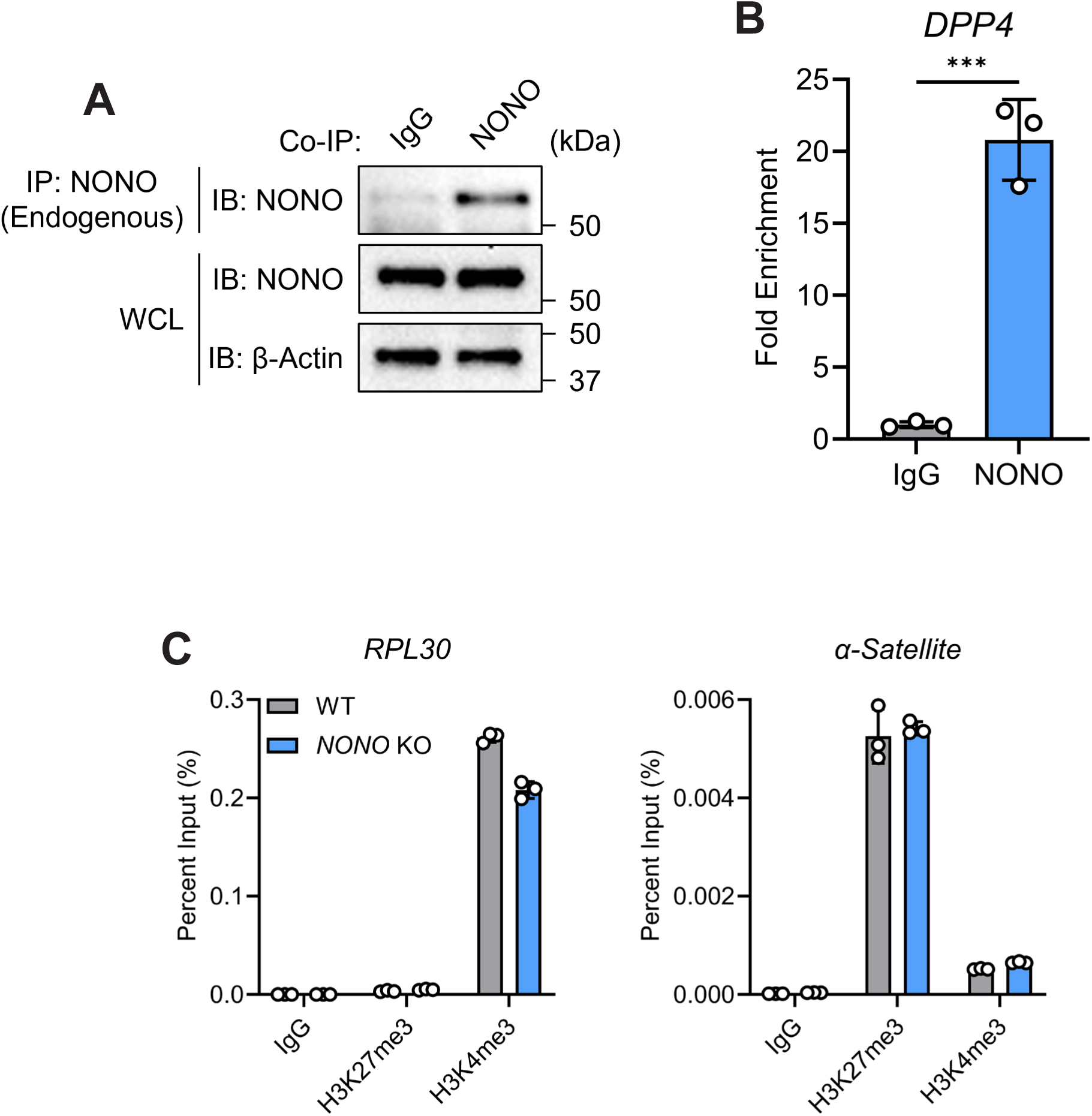
NONO couples chromatin remodeling to expression of the MERS-CoV entry receptor: (A-B) Endogenous NONO RNA immunoprecipitation from WT HAP1 cells. (A) NONO immunoprecipitation efficiency (immunoblot). (B) Expression of NONO-bound *DPP4* (qRT-PCR). (C) ChIP-qPCR showing H3K27me3 and H3K4me3 histone modifications at *RPL30* and *α-Satellite* from WT or *NONO* KO HAP1 cells. Data are expressed as means (n=3) ± SD, ***p < 0.001, (Welch’s t-test) and are representative of 2-3 independent experiments.

## References

1. Flynn, R.A., Belk, J.A., Qi, Y., Yasumoto, Y., Wei, J., Alfajaro, M.M., Shi, Q., Mumbach, M.R., Limaye, A., DeWeirdt, P.C., et al. (2021). Discovery and functional interrogation of SARS-CoV-2 RNA-host protein interactions. Cell 184, 2394–2411.e2316. 10.1016/j.cell.2021.03.012.

2. Lee, S., Lee, Y.-s., Choi, Y., Son, A., Park, Y., Lee, K.-M., Kim, J., Kim, J.-S., and Kim, V.N. (2021). The SARS-CoV-2 RNA interactome. Molecular Cell 81, 2838–2850.e2836. 10.1016/j.molcel.2021.04.022.

3. Labeau, A., Fery-Simonian, L., Lefevre-Utile, A., Pourcelot, M., Bonnet-Madin, L., Soumelis, V., Lotteau, V., Vidalain, P.-O., Amara, A., and Meertens, L. (2022). Characterization and functional interrogation of the SARS-CoV-2 RNA interactome. Cell Reports 39. 10.1016/j.celrep.2022.110744.

4. Hage, A., Janes, M., Shue, B., Markowitz, T.E., Yoon, S., Shannon, J.G., Beare, P.A., Broeckel, R.M., Lack, J.B., Martens, C., and Best, S.M. (2026). The paraspeckle protein NONO potentiates the antiviral innate immune response through chromatin regulation. bioRxiv, 2026.2007.2011.737985. 10.64898/2026.07.11.737985.

5. Ingram, H.B., and Fox, A.H. (2024). Unveiling the intricacies of paraspeckle formation and function. Current Opinion in Cell Biology 90, 102399. 10.1016/j.ceb.2024.102399.

6. Kim, S.J., Ju, J.S., Kang, M.H., Eun, J.W., Kim, Y.H., Raninga, P.V., Khanna, K.K., Győrffy, B., Pack, C.G., Han, H.D., et al. (2020). RNA-binding protein NONO contributes to cancer cell growth and confers drug resistance as a theranostic target in TNBC. Theranostics 10, 7974–7992. 10.7150/thno.45037.

7. Ronchetti, D., Traini, V., Silvestris, I., Fabbiano, G., Passamonti, F., Bolli, N., and Taiana, E. (2024). The pleiotropic nature of NONO, a master regulator of essential biological pathways in cancers. Cancer Gene Therapy 31, 984–994. 10.1038/s41417-024-00763-x.

8. Wei, J., Patil, A., Collings, C.K., Alfajaro, M.M., Liang, Y., Cai, W.L., Strine, M.S., Filler, R.B., DeWeirdt, P.C., Hanna, R.E., et al. (2023). Pharmacological disruption of mSWI/SNF complex activity restricts SARS-CoV-2 infection. Nature Genetics 55, 471–483. 10.1038/s41588-023-01307-z.

9. Chow, R.D., Majety, M., and Chen, S. (2021). The aging transcriptome and cellular landscape of the human lung in relation to SARS-CoV-2. Nature Communications 12, 4. 10.1038/s41467-020-20323-9.

10. Hong, S., Jung, C.H., Han, S., and Park, C.-Y. (2022). Increasing Age Associated with Higher Dipeptidyl Peptidase-4 Inhibition Rate Is a Predictive Factor for Efficacy of Dipeptidyl Peptidase-4 Inhibitors. Diabetes Metab J 46, 63–70. 10.4093/dmj.2020.0253.

11. Deinhardt-Emmer, S., Deshpande, S., Kitazawa, K., Herman, A.B., Bons, J., Rose, J.P., Kumar, P.A., Anerillas, C., Neri, F., Ciotlos, S., et al. (2024). Role of the Senescence-Associated Factor Dipeptidyl Peptidase 4 in the Pathogenesis of SARS-CoV-2 Infection. Aging Dis 15, 1398–1415. 10.14336/ad.2023.0812.

12. Seys, L.J.M., Widagdo, W., Verhamme, F.M., Kleinjan, A., Janssens, W., Joos, G.F., Bracke, K.R., Haagmans, B.L., and Brusselle, G.G. (2018). DPP4, the Middle East Respiratory Syndrome Coronavirus Receptor, is Upregulated in Lungs of Smokers and Chronic Obstructive Pulmonary Disease Patients. Clinical Infectious Diseases 66, 45–53. 10.1093/cid/cix741.

13. Almeida-da-Silva, C.L., Matshik Dakafay, H., Liu, K., and Ojcius, D.M. (2021). Cigarette Smoke Stimulates SARS-CoV-2 Internalization by Activating AhR and Increasing ACE2 Expression in Human Gingival Epithelial Cells. International Journal of Molecular Sciences 22, 7669. 10.3390/ijms22147669.

14. Dighe, A., Jombart, T., Van Kerkhove, M.D., and Ferguson, N. (2019). A systematic review of MERS-CoV seroprevalence and RNA prevalence in dromedary camels: Implications for animal vaccination. Epidemics 29, 100350. 10.1016/j.epidem.2019.100350.

15. Bernard-Stoecklin, S., Nikolay, B., Assiri, A., Bin Saeed, A.A., Ben Embarek, P.K., El Bushra, H., Ki, M., Malik, M.R., Fontanet, A., Cauchemez, S., and Van Kerkhove, M.D. (2019). Comparative Analysis of Eleven Healthcare-Associated Outbreaks of Middle East Respiratory Syndrome Coronavirus (Mers-Cov) from 2015 to 2017. Scientific Reports 9, 7385. 10.1038/s41598-019-43586-9.

16. Killerby, M.E., Biggs, H.M., Midgley, C.M., Gerber, S.I., and Watson, J.T. (2020). Middle East Respiratory Syndrome Coronavirus Transmission. Emerg Infect Dis 26, 191–198. 10.3201/eid2602.190697.

17. Li, K., Wohlford-Lenane, C., Bartlett, J.A., and McCray, P.B. (2022). Inter-individual Variation in Receptor Expression Influences MERS-CoV Infection and Immune Responses in Airway Epithelia. Frontiers in Public Health Volume 9-2021.

18. Algaissi, A., Agrawal, A.S., Han, S., Peng, B.-H., Luo, C., Li, F., Chan, T.-S., Couch, R.B., and Tseng, C.-T.K. (2019). Elevated Human Dipeptidyl Peptidase 4 Expression Reduces the Susceptibility of hDPP4 Transgenic Mice to Middle East Respiratory Syndrome Coronavirus Infection and Disease. The Journal of Infectious Diseases 219, 829–835. 10.1093/infdis/jiy574.

19. Chan, K.K., Tan, T.J.C., Narayanan, K.K., and Procko, E. (2021). An engineered decoy receptor for SARS-CoV-2 broadly binds protein S sequence variants. Science Advances 7, eabf1738. doi:10.1126/sciadv.abf1738.

20. Nishioka, K., Sakai, Y., Motooka, D., Iwata-Yoshikawa, N., Tojo, H., Matoba, S., Nagata, N., Nakaya, T., Arimori, T., and Hoshino, A. (2025). Engineered DPP4 decoy confers broad-spectrum inhibition of MERS-CoV infection. Cell Biomaterials 1. 10.1016/j.celbio.2025.100018.

21. Kathman, S.G., Koo, S.J., Lindsey, G.L., Her, H.-L., Blue, S.M., Li, H., Jaensch, S., Remsberg, J.R., Ahn, K., Yeo, G.W., et al. (2023). Remodeling oncogenic transcriptomes by small molecules targeting NONO. Nature Chemical Biology 19, 825–836. 10.1038/s41589-023-01270-0.

22. Landeras-Bueno, S., Jorba, N., Pérez-Cidoncha, M., and Ortín, J. (2011). The Splicing Factor Proline-Glutamine Rich (SFPQ/PSF) Is Involved in Influenza Virus Transcription. PLOS Pathogens 7, e1002397. 10.1371/journal.ppat.1002397.

23. Murray-Nerger, L.A., Lozano, C., Burton, E.M., Liao, Y., Ungerleider, N.A., Guo, R., and Gewurz, B.E. (2024). The nucleic acid binding protein SFPQ represses EBV lytic reactivation by promoting histone H1 expression. Nature Communications 15, 4156. 10.1038/s41467-024-48333-x.

24. Zhou, B., Wu, F., Han, J., Qi, F., Ni, T., and Qian, F. (2019). Exploitation of nuclear protein SFPQ by the encephalomyocarditis virus to facilitate its replication. Biochemical and Biophysical Research Communications 510, 65–71. 10.1016/j.bbrc.2019.01.032.

25. Zhang, S., Ingram, H., Cooper, J., Naveed, A., Kathman, S.G., Lindsey, G.L., Liu, T., Bond, C.S., Fletcher, J.I., Cravatt, B.F., and Fox, A.H. (2025). NONO Maintains SREBP-Regulated Cholesterol Biosynthesis via RNA Binding in Neuroblastoma. The FASEB Journal 39, e71051. 10.1096/fj.202403267RR.

26. Yuan, S., Chu, H., Chan, J.F.-W., Ye, Z.-W., Wen, L., Yan, B., Lai, P.-M., Tee, K.-M., Huang, J., Chen, D., et al. (2019). SREBP-dependent lipidomic reprogramming as a broad-spectrum antiviral target. Nature Communications 10, 120. 10.1038/s41467-018-08015-x.

27. Theken, K.N., Tang, S.Y., Sengupta, S., and FitzGerald, G.A. (2021). The roles of lipids in SARS-CoV-2 viral replication and the host immune response. Journal of Lipid Research 62, 100129. 10.1016/j.jlr.2021.100129.

28. Soares, V.C., Dias, S.S.G., Santos, J.C., Azevedo-Quintanilha, I.G., Moreira, I.B.G., Sacramento, C.Q., Fintelman-Rodrigues, N., Temerozo, J.R., da Silva, M.A.N., Barreto-Vieira, D.F., et al. (2023). Inhibition of the SREBP pathway prevents SARS-CoV-2 replication and inflammasome activation. Life Science Alliance 6, e202302049. 10.26508/lsa.202302049.

29. Soultsioti, M., de Jong, A.W.M., Blomberg, N., Tas, A., Giera, M., Snijder, E.J., and Bárcena, M. (2025). Perturbation of de novo lipogenesis hinders MERS-CoV assembly and release, but not the biogenesis of viral replication organelles. Journal of Virology 99, e02282–02224. 10.1128/jvi.02282-24.

30. Mitchell Hugh, D., Kyle, J., Engbrecht, K., Berger, M., Oxford Kristie, L., and Sims Amy, C. (2026). Increased triacylglyceride and ceramide levels are key for MERS-CoV replication. mSphere 11, e00523–00525. 10.1128/msphere.00523-25.

31. Martin, M. (2011). Cutadapt removes adapter sequences from high-throughput sequencing reads. EMBnet. journal 17, 10–12.

32. Dobin, A., Davis, C.A., Schlesinger, F., Drenkow, J., Zaleski, C., Jha, S., Batut, P., Chaisson, M., and Gingeras, T.R. (2013). STAR: ultrafast universal RNA-seq aligner. Bioinformatics 29, 15–21. 10.1093/bioinformatics/bts635.

33. Li, B., and Dewey, C.N. (2011). RSEM: accurate transcript quantification from RNA-Seq data with or without a reference genome. BMC Bioinformatics 12, 323. 10.1186/1471-2105-12-323.

34. Law, C.W., Chen, Y., Shi, W., and Smyth, G.K. (2014). voom: precision weights unlock linear model analysis tools for RNA-seq read counts. Genome Biology 15, R29. 10.1186/gb-2014-15-2-r29.

35. Cheng, X., Yan, J., Liu, Y., Wang, J., and Taubert, S. (2021). eVITTA: a web-based visualization and inference toolbox for transcriptome analysis. Nucleic Acids Research 49, W207–W215. 10.1093/nar/gkab366.

36. Shen, S., Park, J.W., Lu, Z.-x., Lin, L., Henry, M.D., Wu, Y.N., Zhou, Q., and Xing, Y. (2014). rMATS: Robust and flexible detection of differential alternative splicing from replicate RNA-Seq data. Proceedings of the National Academy of Sciences 111, E5593–E5601. doi:10.1073/pnas.1419161111.

37. Li, H., and Durbin, R. (2009). Fast and accurate short read alignment with Burrows–Wheeler transform. Bioinformatics 25, 1754–1760. 10.1093/bioinformatics/btp324.

38. Abascal, F., Acosta, R., Addleman, N.J., Adrian, J., Afzal, V., Ai, R., Aken, B., Akiyama, J.A., Jammal, O.A., Amrhein, H., et al. (2020). Expanded encyclopaedias of DNA elements in the human and mouse genomes. Nature 583, 699–710. 10.1038/s41586-020-2493-4.

39. Danecek, P., Bonfield, J.K., Liddle, J., Marshall, J., Ohan, V., Pollard, M.O., Whitwham, A., Keane, T., McCarthy, S.A., Davies, R.M., and Li, H. (2021). Twelve years of SAMtools and BCFtools. GigaScience 10. 10.1093/gigascience/giab008.

40. Ramírez, F., Ryan, D.P., Grüning, B., Bhardwaj, V., Kilpert, F., Richter, A.S., Heyne, S., Dündar, F., and Manke, T. (2016). deepTools2: a next generation web server for deep-sequencing data analysis. Nucleic Acids Research 44, W160–W165. 10.1093/nar/gkw257.

41. Stark, R., and Brown, G. (2011). DiffBind: differential binding analysis of ChIP-Seq peak data. R package version 100, 2–21.

42. Love, M.I., Huber, W., and Anders, S. (2014). Moderated estimation of fold change and dispersion for RNA-seq data with DESeq2. Genome Biology 15, 550. 10.1186/s13059-014-0550-8.

43. Kondili, M., Fust, A., Preussner, J., Kuenne, C., Braun, T., and Looso, M. (2017). UROPA: a tool for Universal RObust Peak Annotation. Scientific Reports 7, 2593. 10.1038/s41598-017-02464-y.

44. Heinz, S., Benner, C., Spann, N., Bertolino, E., Lin, Y.C., Laslo, P., Cheng, J.X., Murre, C., Singh, H., and Glass, C.K. (2010). Simple Combinations of Lineage-Determining Transcription Factors Prime cis-Regulatory Elements Required for Macrophage and B Cell Identities. Molecular Cell 38, 576–589. 10.1016/j.molcel.2010.05.004.

45. López-Muñoz, A.D., Kosik, I., Holly, J., and Yewdell, J.W. (2022). Cell surface SARS-CoV-2 nucleocapsid protein modulates innate and adaptive immunity. Sci Adv 8, eabp9770. 10.1126/sciadv.abp9770.

46. Letko, M., Miazgowicz, K., McMinn, R., Seifert, S.N., Sola, I., Enjuanes, L., Carmody, A., van Doremalen, N., and Munster, V. (2018). Adaptive Evolution of MERS-CoV to Species Variation in DPP4. Cell Reports 24, 1730–1737. 10.1016/j.celrep.2018.07.045.

47. Hage, A., Bharaj, P., van Tol, S., Giraldo, M.I., Gonzalez-Orozco, M., Valerdi, K.M., Warren, A.N., Aguilera-Aguirre, L., Xie, X., Widen, S.G., et al. (2022). The RNA helicase DHX16 recognizes specific viral RNA to trigger RIG-I-dependent innate antiviral immunity. Cell Reports 38. 10.1016/j.celrep.2022.110434.

48. Corman, V.M., Eckerle, I., Bleicker, T., Zaki, A., Landt, O., Eschbach-Bludau, M., van Boheemen, S., Gopal, R., Ballhause, M., Bestebroer, T.M., et al. (2012). Detection of a novel human coronavirus by real-time reverse-transcription polymerase chain reaction. Eurosurveillance 17, 20285. 10.2807/ese.17.39.20285-en.

49. Scobey, T., Yount, B.L., Sims, A.C., Donaldson, E.F., Agnihothram, S.S., Menachery, V.D., Graham, R.L., Swanstrom, J., Bove, P.F., Kim, J.D., et al. (2013). Reverse genetics with a full-length infectious cDNA of the Middle East respiratory syndrome coronavirus. Proceedings of the National Academy of Sciences 110, 16157–16162. 10.1073/pnas.1311542110.

50. Coleman, C.M., and Frieman, M.B. (2015). Growth and Quantification of MERS-CoV Infection. Curr Protoc Microbiol 37, 15e.12.11–19. 10.1002/9780471729259.mc15e02s37.

